# The nucleotide addition cycle of the SARS-CoV-2 polymerase

**DOI:** 10.1101/2021.03.27.437309

**Authors:** Subhas Chandra Bera, Mona Seifert, Robert N. Kirchdoerfer, Pauline van Nies, Yibulayin Wubulikasimu, Salina Quack, Flávia S. Papini, Jamie J. Arnold, Bruno Canard, Craig E. Cameron, Martin Depken, David Dulin

**Affiliations:** Junior Research Group 2, Interdisciplinary Center for Clinical Research, Friedrich-Alexander-University Erlangen-Nürnberg (FAU), Cauerstr. 3, 91058 Erlangen, Germany; Department of Biochemistry and Institute for Molecular Virology, University of Wisconsin-Madison, Madison, WI 53706; Architecture et Fonction des Macromolécules Biologiques, CNRS and Aix-Marseille Université, UMR 7257, Polytech Case 925, 13009 Marseille, France; Department of Microbiology and Immunology, University of North Carolina School of Medicine, Chapel Hill, NC 27599 USA; Department of Bionanoscience, Kavli Institute of Nanoscience, Delft University of Technology, Van der Maasweg 9, 2629 HZ Delft, The Netherlands; Department of Physics and Astronomy, and LaserLaB Amsterdam, Vrije Universiteit Amsterdam, De Boelelaan 1081, 1081 HV, Amsterdam, The Netherlands

**Author notes:** These authors contributed equally to this work (Subhas Chandra Bera, Mona Seifert).

## Abstract

Coronaviruses have evolved elaborate multisubunit machines to replicate and transcribe their genomes. Central to these machines are the RNA-dependent RNA polymerase subunit (nsp12) and its intimately associated cofactors (nsp7 and nsp8). We have used a high-throughput magnetic-tweezers approach to develop a mechanochemical description of this core polymerase. The core polymerase exists in at least three catalytically distinct conformations, one being kinetically consistent with incorporation of incorrect nucleotides. We provide the first evidence that an RdRp uses a thermal ratchet instead of a power stroke to transition from the pre- to post-translocated state. Ultra-stable magnetic tweezers enables the direct observation of coronavirus polymerase deep and long-lived backtrack that are strongly stimulated by secondary structure in the template. The framework presented here elucidates one of the most important structure-dynamics-function relationships in human health today, and will form the grounds for understanding the regulation of this complex.

## Introduction

SARS-CoV-2 is the third zoonotic coronavirus outbreak in less than twenty years, after SARS-CoV-1 and MERS-CoV. To date, SARS-CoV-2 has infected more than 100 millions people, which has led to more than 2.4 millions deaths, with numbers still on the rise. Though several vaccines are now available, we still lack easily administered antiviral drugs to protect non-vaccinated populations against the current or future coronavirus outbreaks (*1*). The ~30 kb long positive single-stranded ((+)ss) RNA genome of coronaviruses encodes many structural and non-structural proteins (nsp). Amongst the latter are the viral proteins constituting the multi-subunits RNA-dependent RNA polymerase (RdRp) responsible for replication and transcription of the viral genome (*2*). This replication/transcription machinery may differ in composition of accessory factors, but likely have the same core (referend to as polymerase from here on): the RdRp subunit (nsp12) and accessory factors (nsp7 and nsp8) (*3, 4*). Because of its central role in the virus life cycle, the coronavirus polymerase represents a major drug target (*5*). Nucleotide analogues, such as Remdesivir, are the only therapeutic option to treat coronavirus infection, and a precise understanding of the nucleotide addition cycle would tremendously help the development of anti-viral drugs.

The nsp12 structure includes the typical features from (+)ssRNA virus RdRps (*6*), with a cupped right hand shape that include palm, finger and thumb subdomains (*3, 4, 7, 8*). The coronavirus polymerase complex replicates and transcribes the viral genome at a high pace, having the highest nucleotide addition rate measured for any RNA polymerase to date, i.e. ~170 nt/s at 37°C (*9–11*). The fidelity of RdRps is on par with DNA polymerases lacking exonuclease activity (*12*). Though RdRp infidelity enhances viral evolution (*13*), the stability of the large coronavirus genome is not as tolerant to mutation as viruses with smaller genomes. Therefore, to further reduce mutational burdern, the coronavirus genome also encodes a 3’-to-5’ exonuclease, nsp14, that proofreads the RNA product (*14–17*). Importantly, nsp14 also confers coronaviruses with protection against many antiviral nucleotide analogues (*5*).

Despite the coronavirus polymerase being a validated therapeutic target, we have little understanding of both its structure-dynamics-function relationships or the mechanochemistry governing the thousands of reiterative cycles of nucleotide addition. Furthermore, we have no knowledge on the mechanisms that underlies the surveillance of the viral genome replication transcription by other viral co-factors. For example, what is the kinetic or biochemical signal that triggers nsp14 to intervene during replication to excise a mismatch? How and how long does nsp14 intervene? Answers to such questions will facilitate the comprehension of what makes a nucleotide analogue potent, e.g. clarifying how does Remdesivir largely elude nsp14 surveillance while ribavirin (another well-established antiviral nucleotide analogue (*18*)) does not (*5*). However, processes such as nucleotide mismatch incorporation in physiological concentration of all NTPs are asynchronous and transient, making their observation difficult for classic ensemble approaches. Single-molecule assays uniquely enable the direct observation of single enzymatic complex, and sheds light on rare – but essential – biochemical events, such as nucleotide mismatch and analogue incorporation (*19*), which are hidden to even the best-resolved bulk assays. These techniques also reveal the subtle interplay between branched kinetic pathways that characterizes polymerase nucleic acid synthesis activity. Single molecule magnetic and optical tweezers have been successfully applied to investigate polymerases mechanochemistry, with near base-pair resolution on thousands of cycles and under any solution condition (*20*). Tweezers also provide the ability to interrogate the translocation mechanism and efficiency in different biomechanical contexts, e.g. elongating on a single-stranded template, through a double-stranded nucleic acid secondary structure, or bypassing roadblocks caused by bound proteins. These techniques have shown a great success in providing the most complete characterization of the mechanochemical cycles in elongation of many model DNA polymerases (*21–26*), cellular RNA polymerases (*27–29*) and viral RdRps (*19, 30–32*).

Using our recently developed single-molecule high-throughput magnetic tweezers assay (*9*), we reveal here the mechanochemical cycle of nucleotide addition by the SARS-CoV-2 polymerase. There is an expectation that the multi-factor assemblies supporting transcription and replication of the coronavirus genome will be as complex as the machineries transcribing and replicating DNA in the human nucleus. The studies of the core polymerase complex reported here represent the first step towards building higher order assemblies and defining the function of each factor. Moreover, our discovery of multiple discrete catalytically-competent forms of the polymerase complex, with only one potentially being surveilled by the proofreading exonuclease, suggest strategies to greatly enhance the potency of “next-generation” anti-coronavirus nucleotide analogues.

## Results

### A high throughput magnetic tweezers assay to investigate SARS-CoV-2 polymerase RNA synthesis kinetics

We investigated the mechanochemistry of the SARS-CoV-2 polymerase during RNA synthesis using a high-throughput magnetic tweezers assay (*9*). We designed an RNA hairpin construct (**Fig. S1A**, **Supplementary Materials**), which provides a 1043 nt ssRNA template when stretched with a force above ~22 pN (**Fig. S1B**). The hairpin construct is flanked at one end by a biotin-labeled handle to attach the magnetic bead, and at the other end by a digoxigenin-labeled handle to anchor the RNA construct to the coverslip glass surface of a flow chamber (**Fig. 1A**). The applied force is controlled by the distance between the magnetic beads and a pair of permanent magnets (*33*). The SARS-CoV-2 polymerase, formed by nsp12, nsp8 and nsp7, assembles at the 3’ end of a ~800 nt long primer and converts the 1043 nt long ssRNA into dsRNA while elongating, which proportionally decreases the construct end-to-end extension (**Fig. 1A**). The observed nanometer-scale change in extension is subsequently converted in nucleotides (Supplementary Materials) (*32*). High-throughput magnetic tweezers enable the observation of hundreds of RNA hairpin tethered magnetic beads (**Fig. S1C**), providing the simultaneous acquisition of dozens of SARS-CoV-2 polymerase elongation traces (**Fig. 1B**), from which we characterized both the final product length and the total replication time (**Fig. 1C**).

**Figure 1:**
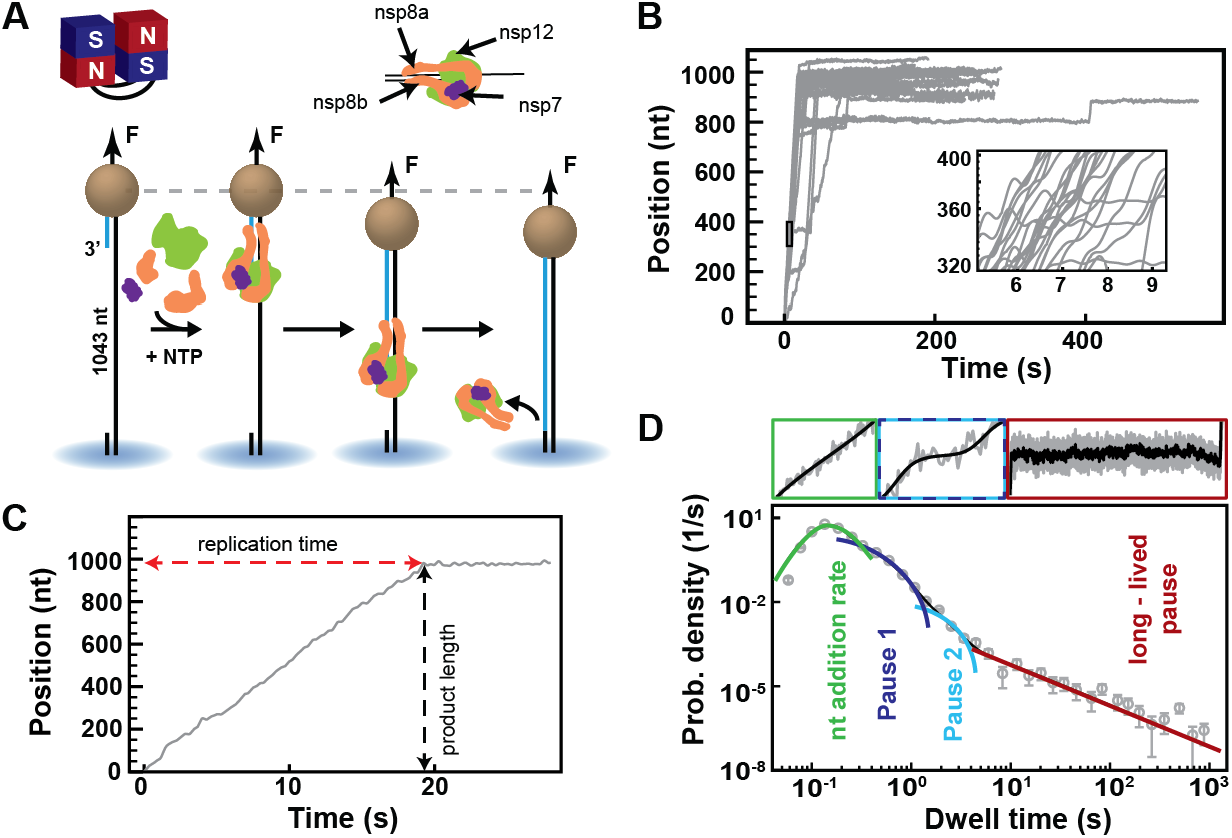
A magnetic tweezers assay to characterize SARS-CoV-2 polymerase elongation kinetics. **(A)** Schematic of the magnetic tweezers assay to monitor RNA synthesis by the SARS-CoV-2 polymerase. A magnetic bead is attached to a glass coverslip surface by an RNA construct including a 1043 nt long ssRNA segment as template, which experiences a constant force F. nsp7, nsp8 and nsp12 assemble at the 3’-end of a ~800 nt long primer ssRNA strand annealed to the template to form an RNA-synthesis competent polymerase. The conversion of the ssRNA template into dsRNA during primer elongation by the polymerase reduces the end-to-end extension of the tether, signaling an elongating polymerase. **(B)** Traces of SARS-CoV-2 RNA synthesis activity acquired at 25 pN, 25 °C and with 500 μM NTP, (inset) showing bursts of nucleotide addition interrupted by pauses of various duration. **(C)** From the SARS-CoV-2 RNA synthesis activity trace the product length (black) and the total replication time (red) are extracted. **(D)** A log-binned histogram of the dwell times extracted from the SARS-CoV-2 polymerase activity traces assembled into a probability density distribution. The above insets are examples of the kinetic events (no pause, short pauses, long pause) that dominate a given dwell time. The distribution has been fitted using a maximum likelihood estimation (MLE) approach using the pause-stochastic model described in the **Materials and Methods** (solid lines). The model includes four different probability distribution functions describing the event that kinetically dominates the dwell time: uninterrupted ten nucleotide additions (green); exponentially distributed Pause 1 and 2 (blue and cyan, respectively); and the power law distributed long-lived pause (red).

As observed with other RNA virus RdRps (*19, 30–32*), the SARS-CoV-2 polymerase activity traces show bursts of nucleotide additions interrupted by pauses of durations ranging from subseconds to hundreds of seconds (**Fig. 1B**). We scanned the activity traces with successive non-overlapping windows of 10 nt, and so measured the duration of ten successive nucleotide addition cycles, coined dwell times, to extract the detailed kinetic information from SARS-CoV-2 polymerase RNA synthesis (**Fig. 1D**).

We previously introduced a stochastic-pausing model to describe RdRp’s dwell time distributions for Φ6, poliovirus, and human rhinovirus C RdRps (*19, 30–32*), and this model was here applied to the SARS-CoV-2 dwell time distributions (**Supplementary Materials**). This model includes four probability distribution functions describing the events that kinetically dominate the dwell time: uninterrupted ten nucleotide additions (gamma distribution), short-lived Pause 1 and Pause 2 (exponential distrinutions), and the long-lived pauses (power law distribution) (**Fig. 1D**). The long-lived pause durations are consistent with a *t*^−3/2^ power-law tail, similar to what is expected from polymerase backtracking (*32, 34*) (**Supplementary Materials**).

Unlike other methods used to analyze polymerase elongation kinetics, the pauses and the pause-free nucleotide addition bursts are characterized without any time binning, using a maximum likelihood estimation (MLE) algorithm applied directly to the dwell times (*32*) (**Supplementary Materials**). Our approach reduces both biases introduced by using arbitrary thresholds to discriminate between pauses and nucleotide addition bursts or by fitting binned data. The model contains seven free parameters to describe SARS-CoV-2 RdRp elongation kinetics over four orders of magnitude in time. This is on par with other models describing the elongation kinetics of the bacterial RNA polymerase (*27, 28, 35*) and replicative DNA polymerases (*24, 25*) at the single-molecule level.

### SARS-CoV-2 polymerase is a processive RNA polymerase

We first investigated whether the observed pauses were related to the exchange of the polymerase itself or its factors. If one of the pauses originated from the exchange of polymerase factor(s), we expect the kinetics of this specific pause to be affected by varying the concentration of proteins in the reaction buffer. To test this, we performed two types of experiments. In the first set of experiments, we flushed the flow chamber with reaction buffer containing NTP and the polymerase factors at different concentrations of nsp12, i.e. 0.1, 0.2 and 0.4 μM; while maintaining the nsp12:nsp8:nsp7 stoichiometry at 1:9:9, i.e. with an excess of nsp7 and nsp8. In this case, elongation starts as soon as the polymerase has assembled on the primer-template, while the polymerase factors are present in the solution to enable protein exchange. In the second set of experiments, we pre-assembled (PA) the polymerase by incubating the polymerase factors in the flow chamber without NTP, subsequently rinsed the flow chamber to remove any free proteins, and started the polymerase RNA synthesis activity by injecting a reaction buffer solution containing 500 μM NTP (**Supplementary Materials**, **Fig. 2A**).

**Figure 2:**
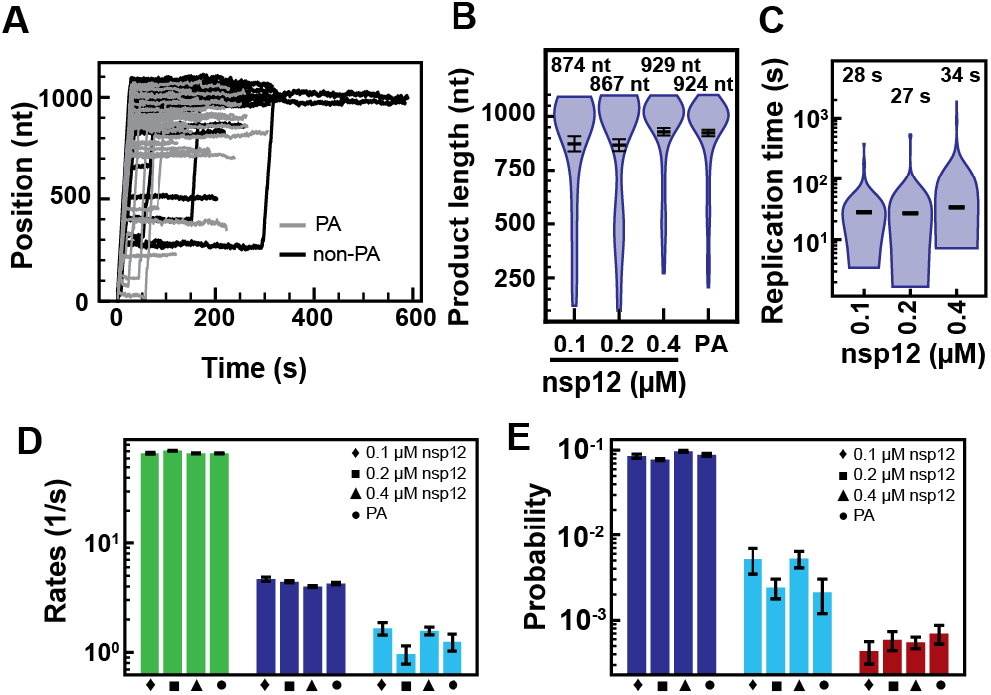
SARS-CoV-2 elongation dynamics does results from polymerase co-factors disassembly or exchange. All experiments were conducted at 35 pN and 25 °C with 500 μM all NTP. (A) Traces of SARS-CoV-2 RNA synthesis activity of pre-assembled (PA, grey) and non-pre-assembled (non-PA, black) polymerase. PA polymerase may start synthesizing RNA while flushing reaction buffer with NTP, and therefore only the part of trace collected at the end of the infusion is represented, demonstrating shorter traces. (B) Replication time and (C) product length of SARS-CoV-2 polymerase as a function of nsp12, nsp7 and nsp8 concentration at a constant stoichiometry of 1:9:9. The median replication times and the mean product lengths are indicated above the violin plots and represented as horizontal thick black lines, flanked by one standard deviation error bars extracted from 1000 bootstraps. (D) Nucleotide addition rates (green bars), Pause 1 (dark blue bars) and Pause 2 (cyan bars) exit rates for PA polymerase and concentration of nsp12 as described in the panels and nsp7 and nsp8 concentrations as described in (B, C). (E) Probabilities to enter Pause 1 (dark blue bars), Pause 2 (cyan bars) and long-lived pauses (red bars) for the conditions described in (D). Error bars are one standard deviation extracted from 100 bootstraps.

In the presence of the polymerase factors in solution, the product length and replication time were not significantly changed for the different polymerase factors concentrations we evaluated, with mean values of (899 ± 28) nt and (30 ± 3) s, respectively (**Fig. 2BC**). A dwell time analysis of these traces similarly reported that the nucleotide addition, Pause 1, Pause 2, and the long-lived pause distributions were also unresponsive to changes in concentration of the polymerase factors in the flow cell (**Fig. 2DE**, **Fig. S2A**). We then evaluated the elongation kinetics of the pre-assembled complex. Because the traces may have started at an undefined time during the injection of the reaction buffer containing NTPs, we did not evaluate the replication time for the preassembled polymerase activity traces. We performed a dwell-time analysis of these traces (**Fig. S2B**), which showed again that none of the parameters of the stochastic-pause model were affected by the absence of viral proteins in the solution. We conclude that neither the product length, the replication time, nor the dynamics observed in the traces, resulted from viral protein disassembly or exchange during RNA synthesis. The coronavirus polymerase is a processive RNA polymerase.

### The short pauses are the signature of slow nucleotide addition pathways distinct from the nucleotide addition burst pathway

We subsequently investigated how nucleotide concentration affects the RNA synthesis kinetics to determine which state is catalytically competent, the saturating NTP concentration of such states, and their nucleotide addition kinetics parameters, such as the Michaelis-Menten (MM) constant *K_M_* and the maximum nucleotide-addition rate *k_max_*. We varied the NTP concentration between 10 μM and 1 mM at a constant force of 25 pN (**Fig. 3A**). We noted that decreasing the NTP concentration significantly increases the average total replication time from (21 ± 1) s to (283 ± 9) s (**Fig. 3B**), without affecting the product length (**Fig. S3A**). The shape of the dwell-time distribution is also affected by decreasing the concentration of NTP: Pause 1 and Pause 2 shoulders inflate dramatically, while the nucleotide addition peak fades away behind the Pause 1 distribution (**Fig. 3C**, **Fig. S3B**). Fitting the distributions using the stochastic-pausing model showed that SARS-CoV-2 polymerase nucleotide addition rate is constant at ~76 nt · s^−1^) over the whole range of NTP concentration (**Fig. 3D**). This is a rather surprising result, as one would expect polymerase nucleotide addition rate to increase with NTP concentration, up to some maximal rate *k_max_* when NTP binding is saturated. The observed NTP concentration independence suggests that the chemistry of nucleotide addition is not rate limiting, and must be followed by a second (nearly) irreversible step that dominates the overall reaction timescale within the NTP concentration explored. Another surprising result is that Pause 1 and Pause 2 exit rates decreased by one order of magnitude when decreasing in NTP concentration (**Fig. 3D**, **Table S1**). This indicates that Pause 1 and Pause 2 are the kinetic signatures of a slow and a very slow nucleotide additions (SNA and VSNA) pathways, in addition to the nucleotide addition bursts (NAB) pathway. Pause 1 and Pause 2 exit rates as a function of NTP concentrations are well described by MM kinetics (**Fig. 3D)**, with the exit rate written as

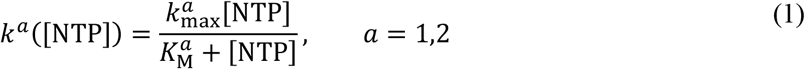

**Figure 3:**
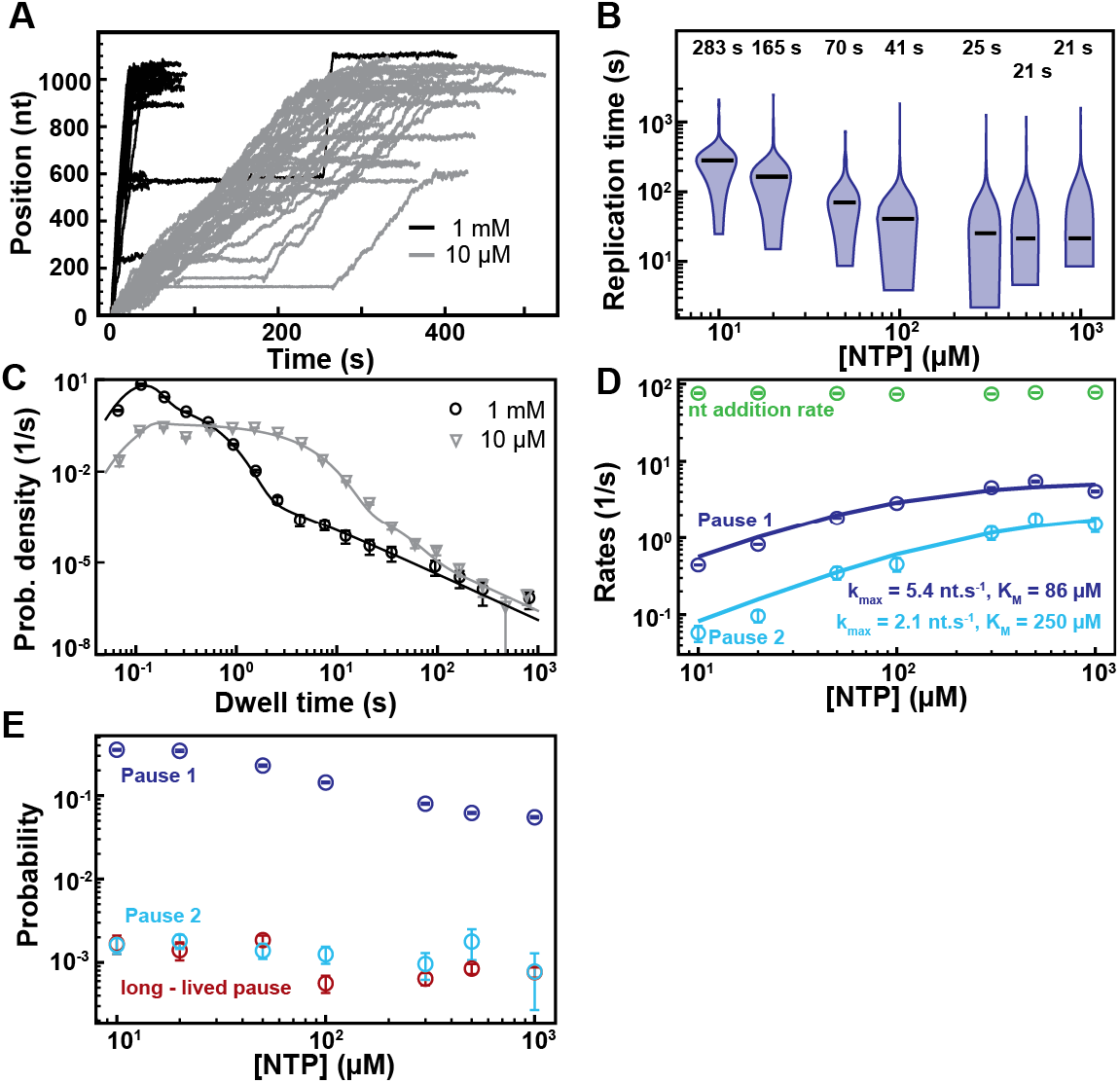
Pause 1 and Pause 2 are the kinetic signatures of low efficiency catalytic states. All experiments were conducted at 25 pN, 25 °C, 0.6 μM nsp12, 1.8 μM nsp7 and 1.8 μM nsp8. **(A)** SARS-CoV-2 polymerase activity traces at 1 mM NTP (black) and 10 μM NTP (grey). **(B)** The replication time as a function of NTP concentration. The median replication times are indicated above the violin plots and represented as horizontal thick black lines, flanked by one standard deviation error bars extracted from 1000 bootstraps. **(C)** The dwell time distributions and their corresponding MLE fits (solid lines) at 1 mM (black circles) and 10 μM NTP (grey triangles). **(D)** The nucleotide addition rates (green circle), Pause 1 (dark blue circle) and Pause 2 (cyan circle) exit rates as a function of NTP concentration. The solid lines are Michaelis-Menten fit to Pause 1 and Pause 2 exit rates NTP concentration dependence. The extracted Michaelis-Menten parameters are indicated next to the fits using the color code described above. **(E)** The probabilities for Pause 1 (dark blue circle), Pause 2 (cyan circle) and long-lived pauses (red circle). The error bars in (C) denote one standard deviation extracted from 1000 bootstraps. The error bars in (D, E) are one standard deviation extracted from 100 bootstraps.

where 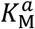 is a force and translocation dependent analog of the MM constant, and 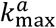 is the maximal exit rate from Pause *a*. We extracted 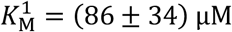 and 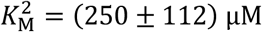, and 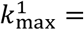 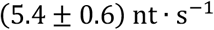 and 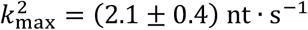 at a force of 25 pN (**Fig. 3D**). Pause 1 probability was constant down to 500 μM NTP, and subsequently increased by almost 6-fold when further reducing NTP concentration down to 20 μM (**Fig. 3E**, **Table S1**). We note that Pause 1 probability did not increase further below 20 μM NTP, which is likely due to our analysis being unable to fit the gamma distributions at large Pause 1 probability (above ~0.3) (**Fig. S3B**). Pause 2 probability increased by ~2.3-fold (**Fig. 3E**, **Fig. S3C**), i.e. not proportionally to Pause 1. The change in probability with NTP concentration indicates that SNA and VSNA pathways are distinct from the NAB pathway. To verify whether the kinetics response of the polymerase to the NTP titration was not just specific to the 25 pN force, we performed the same set of experiments at 35 pN (**Fig. S4A**). The trend observed at 25 pN was conserved at 35 pN. The median replication time decreased when increasing NTP concentration (**Fig. S4B**), while the average product length remained constant (**Fig. S4C**). The nucleotide addition rate was constant for NTP concentrations down to 50 μM NTP, and then decreased, which likely resulted from the poor MLE fit of the gamma-distribution due to the large Pause 1 probability (**Fig. S4DE**). Pause 1 and Pause 2 exit rates remained well described by the effective MM equation (**Equation 1**), i.e. 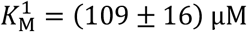 and 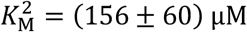, and 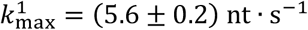 and 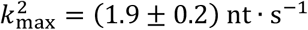 (**Fig. S4E**). Pause 1 and Pause 2 probability increased when decreasing NTP concentration, but faster than at 25 pN suggesting a force dependence for these pauses (**Fig. S4F**, **Table S1**). In conclusion, the coronavirus polymerase presents three catalytic pathways, where Pause 1 and Pause 2 are respectively the kinetic signatures of a slow and a very-slow-nucleotide addition pathways branching off from the nucleotide addition burst pathway (described by the nucleotide addition rate), and the long-lived pauses are catalytically incompetent.

### A large conformational change dominates the coronavirus polymerase nucleotide addition cycle kinetics

By varying the tension on the template, we probed whether a rate-limiting conformational change occurs during the nucleotide addition cycle (*36*). Modelling the experimental data enables us to extract the magnitude of the conformational change along the DNA, as well as the conformer-conversion rate at zero force. Increasing the force from 20 to 60 pN increased the total duration of the activity traces and the number of pauses in these traces (**Fig. 4A**). The median replication time increased by 4-fold, whereas the average product length only mildly decreased (**Fig. 4BC**). Surprisingly, even at forces as high as 60 pN, the SARS-CoV-2 polymerase demonstrated a strong RNA synthesis activity (**Fig. 4A**).

**Figure 4:**
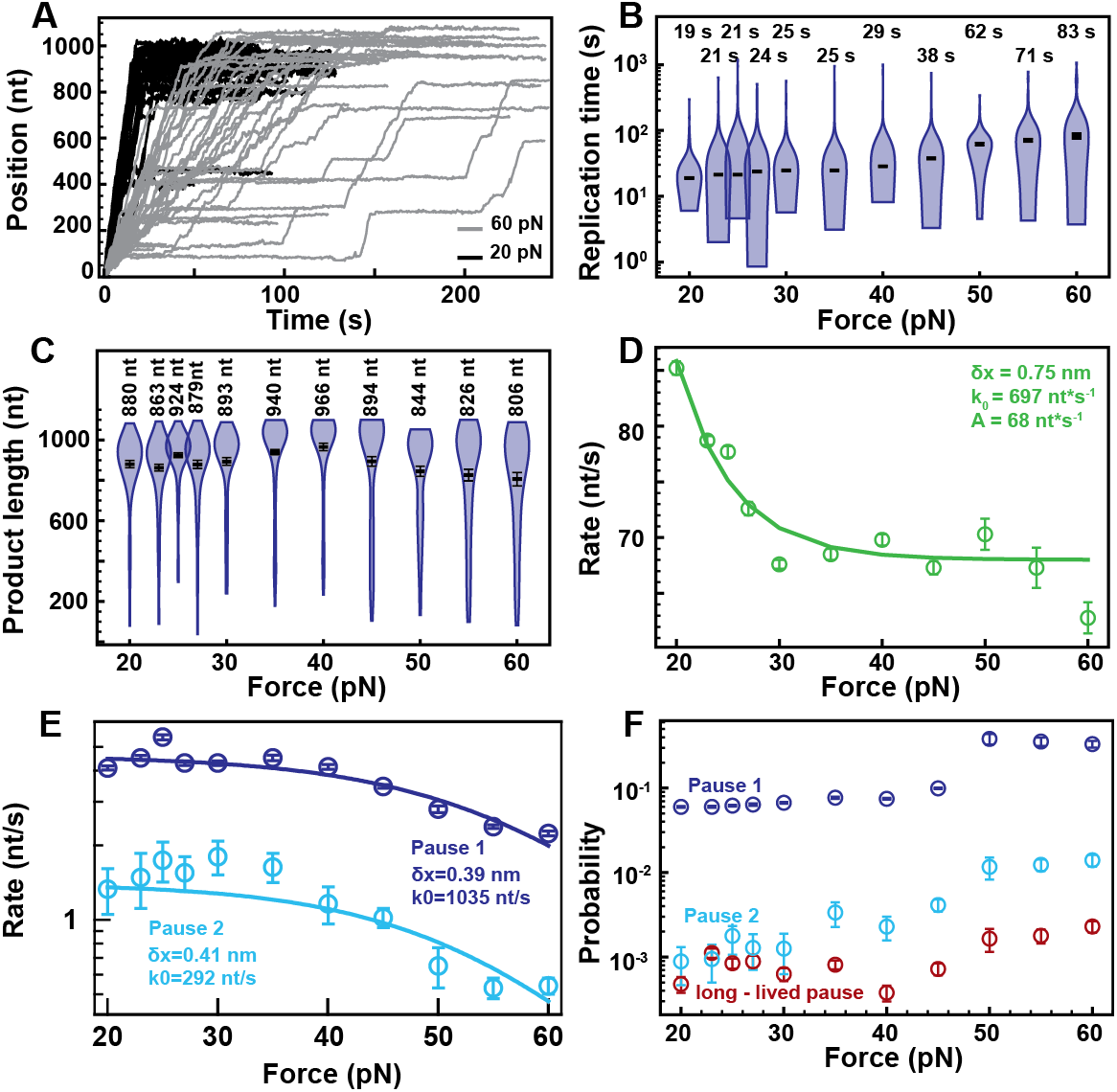
The force dependence of SARS-CoV-2 polymerase kinetics indicates a rate-limiting large conformational change. All experiments were conducted at 500 μM NTP, 25 °C, 0.6 μM nsp12, 1.8 μM nsp7 and 1.8 μM nsp8. **(A)** SARS-CoV-2 polymerase activity traces at 20 pN (black) and 60 pN (grey). **(B)** Replication time and **(C)** product length of SARS-CoV-2 polymerase as a function of force. The median replication times and the mean product lengths are indicated above the violin plots and represented as horizontal thick black lines, flanked by one standard deviation error bars extracted from 1000 bootstraps. **(D)** The nucleotide addition rate of elongating SARS-CoV-2 polymerase as a function of force. The data were fitted (solid line) with the Arrhenius equation (**Equation 2**). **(E)** Pause 1 (dark blue circle) and Pause 2 (cyan circle) exit rates as a function of force. The dashed lines are the fits of the corresponding rates using **Equation (3)**. **(F)** The probabilities to enter Pause 1 (dark blue circle), Pause 2 (cyan circle) and long-lived pause (red circle) as a function of force. The error bars in (D-F) are one standard deviation extracted from 100 bootstraps.

We used the stochastic-pausing model (**Supplementary Materials**, **Fig. 1D**) to fit the dwell-time distributions obtained from the activity traces collected at forces varying from 20 pN to 60 pN (**Fig. S5A-J**). The nucleotide addition rate decreased exponentially between 20 pN and 30 pN, and remained mostly constant for larger forces (**Fig. 4D**, **Table S1**). A similar plateau at high force has been reported for T7 DNA polymerase (*25*). To describe the force dependence of the nucleotide addition rate when using a ssRNA template, we used an Arrhenius equation with an offset rate *A*

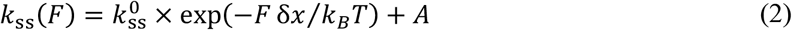

where 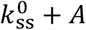 is the nucleotide addition rate at zero force, *F* is the applied force, δ*x* is the distance to the transition state of the reaction, and *k_B_T* is the thermal energy. We extracted *A* = (68 ± 1) nt · s^−1^ and 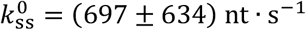, and consequently also a very high nucleotide addition rate on par with recent estimations (*10, 11*). The distance to the transition state δ*x* = (0.75 ± 0.18) nm is larger than a single base distance, which may indicate a large polymerase conformational change not due to translocation.

Pause 1 and Pause 2 kinetics were also force dependent. Their respective exit rates were constant at tensions up to ~35 pN, and decreased at higher forces (**Fig. 4E**, **Table S1**). Our data are consistent with Pause 1 and Pause 2 exiting through at least two irreversible kinetic steps: one NTP addition step that is rate limiting at low forces (up to ~35 pN), with a relatively weak force dependence, and another one strongly force dependent related to a conformational rearrangement that is rate limiting at high forces. We see no appreciable force dependence in the low force regime when the polymerase elongates on a ssRNA template and we therefore described the observed average pause-exit rate as

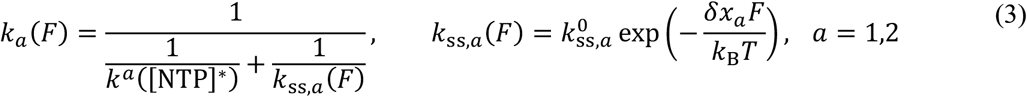

where *k^a^*([NTP]*) parameters are extracted from the fit of **Equation (1)** to Pause 1 and Pause 2 exit rates as a function of NTP concentration at 25 pN (**Fig. 3D**), i.e. in the weakly force dependent regime (**Fig. 4E**), using [NTP]* = 500 μM. **Equation (3)** described well the observed Pause 1 and Pause 2 exit rates, and we extracted 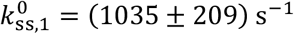 and δ*x*_1_ = (0.39 ± 0.02) nm for Pause 1, and 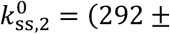 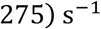 and δ*x*_2_ = (0.41 ± 0.07) nm for Pause 2 (**Fig. 4E**). The probability of Pause 1 and Pause 2 also increased when raising the force from 20 to 45 pN by almost 2- and 5-fold, respectively (**Fig. 4F, Fig. S5K**, **Table S1**), which indicates that the polymerase branches off from the NAB to the SNA and the VSNA pathways from a pre-translocated state that is further populated with increasing tension. The long-lived pause probability showed no force dependence up to 45 pN (**Fig. 4F**). At 50 pN, all pauses showed a ~3-fold jump in probability, which may originate from a change in the dsRNA helix conformation from A- to S-form when being overstretched (*37, 38*).

In conclusion, the second irreversible and rate-limiting step of the nucleotide addition cycle is force-dependent and related to a conformational change more than 3-times larger than a single base translocation. The two low-efficiency catalytic pathways (Pause 1 and Pause 2) also support a two successive irreversible steps model, where the first one is NTP concentration dependent and the second one is force dependent.

### Duplex RNA barrier to the polymerase translocation increases Pause 1 probability and induces polymerase backtracking

The coronavirus genome is heavily structured (*39*), and it is therefore particularly relevant to interrogate the ability of SARS-CoV-2 polymerase to elongate through a closed RNA hairpin (**Fig. 5A**). The conversion of the hairpin into a dsRNA tether happens in two phases. First, the polymerase progresses by unwinding the hairpin stem until its end. Second, the polymerase converts the remaining single stranded RNA into a dsRNA. The first part results in an increase of the tether length by one base and the difference between one base and one base pair for every nucleotide addition cycle. The second part results from the conversion of the ssRNA template into a dsRNA product, as described in **Fig. 1A**. This leads to either a decrease or an increase in extension for forces above and below ~10 pN, respectively (traces at 20 pN and 9 pN in **Fig. 5B**) (*32*). Here, we focused our investigation only on the initial ~0.5 kbp hairpin unwinding activity of the polymerase. At the end of each trace, we verified whether the polymerase fully converted the template into a linear dsRNA by increasing the force to 45 pN, as dsRNA extension is rather constant in this force range, unlike ssRNA (**Fig. S6AB**) (*32*). We discarded the traces that showed a large difference in extension (**Fig. S6B**). A direct comparison of the polymerase activity traces at either 9 pN or 20 pN show that unwinding the hairpin stem at low assisting force induced an increase in the number of pauses and their duration (**Fig. 5B**).

**Figure 5:**
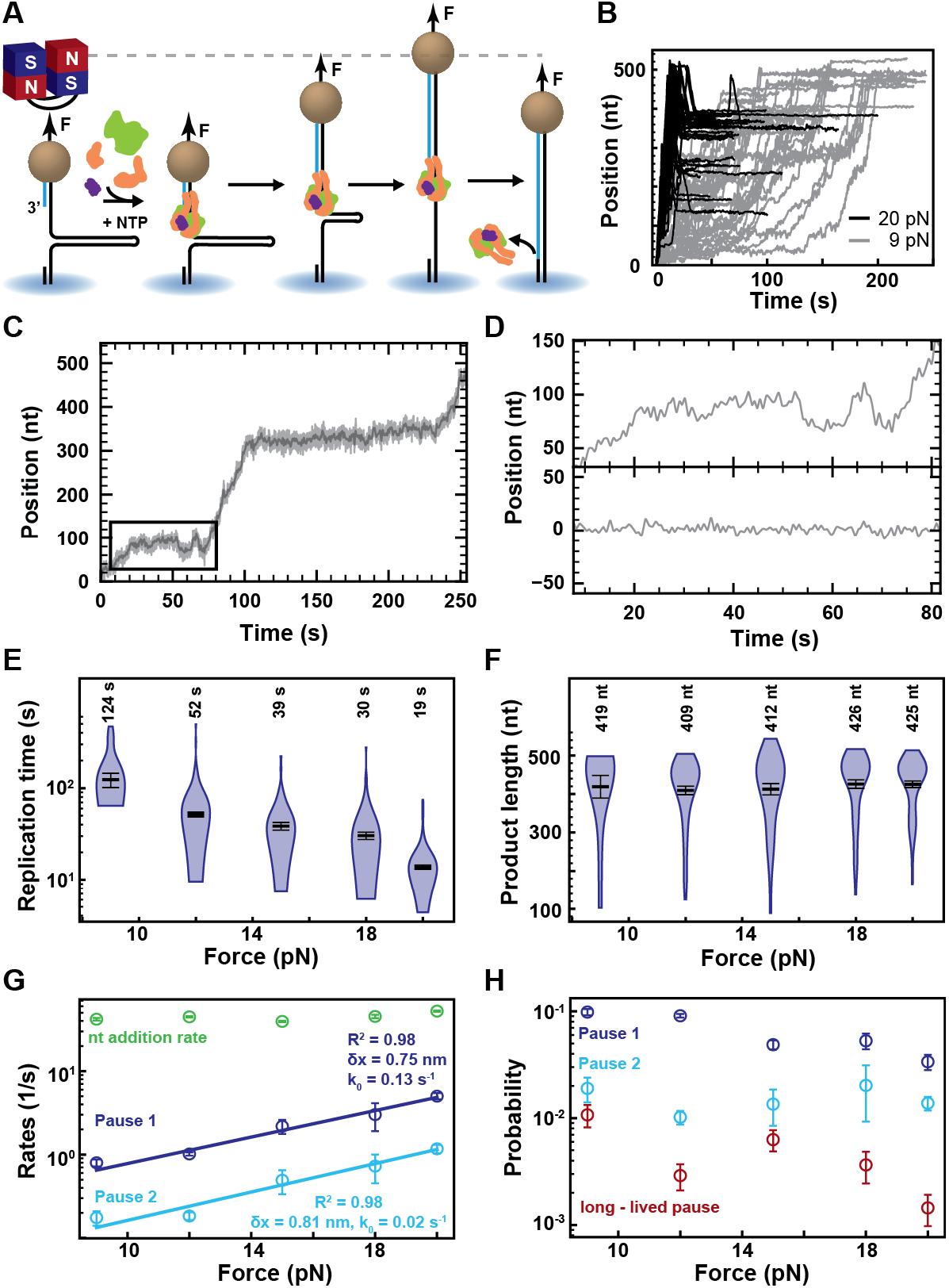
Template secondary structures represent a strong barrier to the elongating SARS-CoV-2 polymerase. All experiments were conducted at 500 μM NTPs, 25 °C, 0.2 μM nsp12, 1.8 μM nsp7 and 1.8 μM nsp8. **(A)** The same hairpin construct is used here as in Figure 1A, though at forces lower than the opening force and the polymerase has to unwind the 499 nt hairpin stem to elongate the primer, leading to a net increase in the extension of end-to-end tether extension of 1 bp + 1 nt until reaching the end of the stem. From the end of the hairpin stem, the change in tether extension is described by the difference in extension of a ssRNA and dsRNA as a function of force, as described in Figure 1A. **(B)** SARS-CoV-2 polymerase activity traces on dsRNA at 20 pN (black) and 9 pN (grey). At 20 pN, an increase in extension appears in the first part of the trace when the polymerase unwinds the hairpin stem, and a subsequent decrease ends the activity trace in the second part, which was converted as the first part of the trace and therefore the values do not reflect the actual number of incorporated nucleotides. We consider only the first part for subsequent analysis. **(C)** SARS-CoV-2 polymerase activity trace at 9 pN using the ultra-stable magnetic tweezers in 100x, 500 μM NTP acquired at 58 Hz (grey), low-pass filtered at 1 Hz (dark grey). (D) Top, zoom-in of the low-pass filtered trace in the black box in (C). Bottom, 1 Hz low-pass filtered trace of a tether without polymerase activity acquired simultaneously. **(E)** Replication time and **(F)** product length of SARS-CoV-2 polymerase on dsRNA as a function of force. The median replication times and the mean product lengths are indicated above the violin plots and represented as horizontal thick black lines, flanked by one standard deviation error bars extracted from 1000 bootstraps. **(G)** The nucleotide addition (green circle), Pause 1 (dark blue circle) and Pause 2 (cyan circle) exit rates as a function of force. The solid lines represent Arrhenius fits to Pause 1 and Pause 2 exit rates (**Equation 1**) **(H)** Probabilities to enter Pause 1 (dark blue circle), Pause 2 (cyan circle) and long-lived pauses (red circle) as a function of force. The error bars in (E, F) are one standard deviation extracted from 100 bootstraps.

Furthermore, we observed an increase in the magnetic bead position fluctuation during the long-lived pauses at 9 pN. To determine whether these fluctuations originated from the polymerase behavior or experimental noise, we repeated the experiments at 9 pN to further improve resolution and stability. To this end, we increased the objective magnification by 2-fold to 100x to reduce the tracking noise (*40*). Furthermore, we introduced a custom autofocus, which significantly increased the stability of the measurement (**Fig. S7A-E**, **Supplementary Materials**), with resolution not being impacted by drift for a duration as long as ~100 s, i.e. a ~50-fold improvement compared to previous studies in similar conditions (*40–42*) (**Supplementary Materials**, **Fig. S7DE**). Using these improved experimental conditions, we were able to confirm that the fluctuations observed in **Fig. 5B** originated from the polymerase position fluctuation (**Fig. 5CD, Fig. S8**). Our data show that the coronavirus polymerase may backtrack as far as ~30 nt upstream the last incorporated nucleotide (**Fig. 5CD, Fig. S8**). Our results are the first direct observation of SARS-CoV-2 polymerase backtracks, and support a recent structural study on a preassembled backtracked coronavirus polymerase (*43*).

Decreasing the force from 20 pN to 9 pN increased the median replication time by almost an order of magnitude (**Fig. 5E**), while the mean product length remained largely constant (**Fig. 5F**). Applying our stochastic-pausing model to the dwell time distributions (**Fig. S6C**), we observed that the nucleotide addition rate decreased when RNA synthesis is performed against a dsRNA barrier, from (85 ± 1) nt · s^−1^) at 20 pN and ssRNA template to (52 ± 1) nt · s^−1^) at the same force but using a dsRNA template (**Fig. 4DE**). The nucleotide addition rate further decreased to (45 ± 3) nt · s^−1^) at 18 pN to remain largely constant at lower forces (**Fig. 5G**, **Table S1**).

While Pause 1 and Pause 2 exit rates measured at 20 pN were comparable when using either the closed or the open hairpin, lowering the force to 9 pN drastically decreased Pause 1 and Pause 2 exit rates by 6− and 7-fold, respectively (**Fig. 5G**, **Table S1**). The decrease is well described by an Arrhenius equation (**Equation 2**, *A* = 0), providing δ_*x*1,ds_ = (0.75 ± 0.08) nm, δ*x*_2,ds_ = (0.81 ± 0.08) nm, 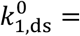 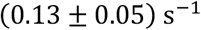 and 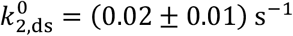. The distance to the transition state for both pauses are similar and consistent with the gain in extension from a single base pair melting, i.e. two single nucleotides (*44*). We therefore suggest that δ*x*_1/2_ relates to single base pair polymerase forward translocation, and therefore translocation becomes rate-limiting for Pause 1 and Pause 2 when the polymerase is elongating through a dsRNA barrier (closed hairpin). The pauses exit rates at zero force, 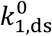 and 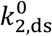, highlight the difficulty for the polymerase alone to exit Pause 1 and Pause 2 when elongating through dsRNA. Finally, elongating through a dsRNA barrier significantly increases the probability to pause: Pause 1, Pause 2 and backtrack pause probabilities increased by 3-, 1.4- and 8-fold while decreasing the force from 20 to 9 pN when using a dsRNA hairpin template (**Fig. 5H**, **Table S1**).

In conclusion, the coronavirus polymerase is rather inefficient at elongating through a duplex RNA template, which suggests that the nsp13 helicase or other viral co-factors assist the polymerase during viral genome replication and transcription.

### A nucleotide addition cycle model for the coronavirus polymerase

To describe the NTP and force dependence of the coronavirus polymerase on either ss- or dsRNA template, we introduce a novel model for the nucleotide addition cycle of the coronavirus polymerase, which may be applicable to other related viral RdRps by extension (**Fig. 6A**, **Fig. S9**). Our model supports a polymerase translocation mechanism at the beginning of the nucleotide addition cycle, where the step forward is thermally activated. This step could be mistaken for a power-stroke as it does not demonstrate a force dependence in rate. The forward translocation is then stabilized by NTP binding, followed by the closure of the active site (*45*). Pause 1 probability increases when raising the applied tension on the ssRNA template (open hairpin) or decreasing the applied tension on the dsRNA template (closed hairpin). We therefore conclude that the SNA pathway (Pause 1) branches out from the pre-translocated state of the NAB pathway (**Fig. 6A**, **Fig. S9**). Furthermore, Pause 1 probability never becomes null at saturating NTP, but rather plateaus above 500 μM. This indicates that translocation can not be considered equilibrated with respect to NTP binding and Pause 1 entry rate, and is even rate-limiting for both Pause 1 and Pause 2 when using a dsRNA hairpin template. The NTP concentration independence of the SARS-CoV-2 polymerase nucleotide addition rate (NAB pathway) at two different forces (25 and 35 pN) indicates the existance of another irreversible and rate-limiting step following the nucleotide addition chemistry. The force dependence of this state suggests a large conformational change of the polymerase at the end of the nucleotide addition cycle. The SNA and VSNA pathways (Pause 1 and Pause 2) exit rates are also consistent with two successive irreversible steps, one related to the chemistry of nucleotide addition, and the other related to a tension-dependent conformational change. An irreversible conformational change was also observed in a pre-steady state kinetic study of poliovirus RdRp (*12*), and was inferred to be the translocation of the polymerase. Our data however show that the distance to the transition state of the reaction is three times as large as a single base translocation for both the NAB, SNA and VSNA. We therefore conclude that this conformational change cannot be soley due to translocation, and likely represents a large conformational change in the elongating polymerase.

**Figure 6:**
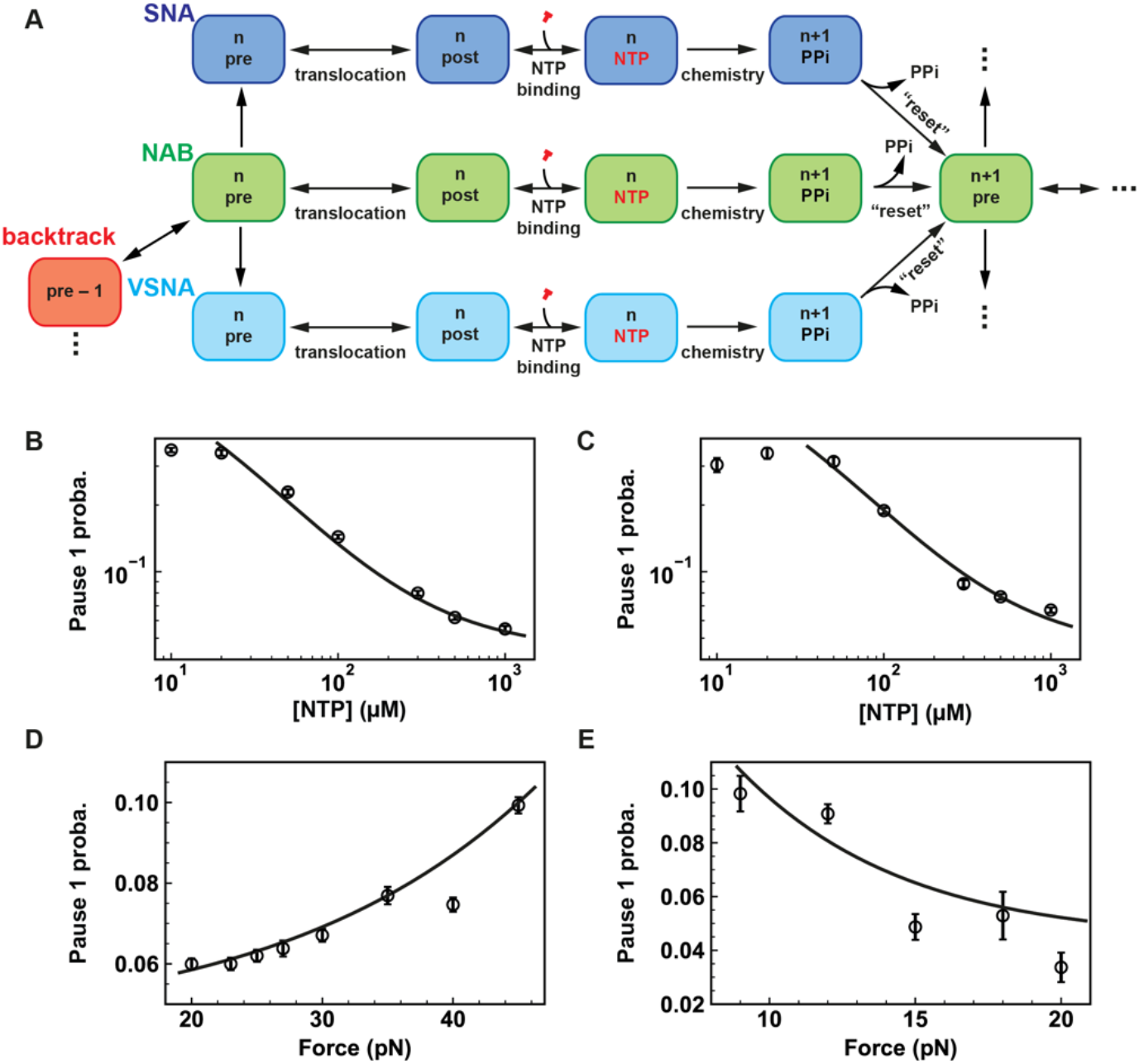
A Mechanochemical model of the coronavirus polymerase nucleotide addition cycle. **(A)** The mechanochemical cycle starts from the pre-translocated state (pre) at position n on the product RNA of the nucleotide addition burst (NAB) pathway. From this position, the polymerase either use thermal energy to translocate forwards onto the post-translocated state, and carry on with nucleotide binding, chemistry (n+1 RNA product length) and a large conformational change that “resets” the polymerase towards the next nucleotide addition cycle, or enter the slow-nucleotide addition (Pause 1) or very-slow-nucleotide addition (Pause 2) (SNA and VSNA, respectively) pathways. The polymerase may also enter a backtrack state, which appears as in the trace as polymerase position behind the maximum position reached on the template and/or long-lived pauses, and is favored by elongating through a closed hairpin. **(B, C)** Pause 1 probabilities as a function of the NTP concentration at either 25 pN or at 35 pN, and **(D, E)** as a function of force for either a ssRNA or dsRNA templates at 500 μM NTP, were fitted with **Equation S4** of the kinetic model describing the entry in Pause 1 (LEC) (**Supplementary Materials**).

Performing a gobal fit of the probability to enter Pause 1 as a function of NTP concentration (both at 25 and 35 pN), and of the applied tension (at 500 μM NTP) (**Supplementary Materials**), we show that our model describes the kinetics of the elongating SARS-CoV-2 polymerase well (**Fig. 6BCD**, **Supplementary Materials**). This model extract a step size from the pre- to the post-translocated states of 0.23 nm, which is in excellent agreement with a single nucleotide translocation distance (**Supplementary Materials**). The model also accounts for the increase in Pause 1 probability as a function of the applied tension when using a dsRNA template (**Fig. 6E**, **Supplementary Materials**). The long-lived pauses are the kinetic signature of polymerase backtracking, which dramatically increased when the polymerase elongates through a dsRNA hairpin. We conclude that the backtrack branches off from the NAB pathway pre-translocated state (**Fig. 6A**, **Fig. S7**).

## Discussion

Understanding the mechanism, regulation, and inhibition of the SARS CoV-2 replication and transcription complex necessitates a precise and complete description of the nucleotide-addition cycle. While structural information has been gathered at an incredible pace (*4, 7, 8, 46*), a mechanistic framework to interpret the structural information is not available. Here, we have interrogated the nucleotide-addition cycle of the SARS CoV-2 core polymerase complex on a one kilobase long template using a high-througput magnetic tweezers approach (*9*). Consistent with previous studies of other RdRps (*19, 30–32*), RNA synthesis by this complex is described best as a series of bursts of nucleotide addition interrupted by pauses of various durations (**Fig. 1**). We show here that these pauses originate from distinct structural/conformational states of the complex and/or transactions performed by the complex, potentially resulting from nucleotide misincorporation. Once initiated and over the 1000 cycles of nucleotide addition monitored, there was no evidence for exchange of the protein components of the complex or dissociation of the complex (**Fig. 2**). These results demonstrate that the SARS-CoV-2 polymerase complex is both stable and processive.

We observed that the nucleotide addition rate of the SARS-CoV-2 polymerase decreased when increasing the tension, and eventually plateaued above 30 pN force at a minimum of ~70 nt/s (**Fig. 4D**). Why does not the polymerase nucleotide addition rate decrease to zero, as observed for φ29 DNA polymerase (*24*)? We suggest that remodeling of the primer-template-polymerase interactions upon changes in tether tension is limited. To verify this hypothesis, we modeled the downstream position of the template strand by aligning the recent cryo-EM structure of an elongating SARS-CoV-2 polymerase (*4*) to the crytal structure of an elongating poliovirus RdRp (*6*). The upstream duplex RNA product aligns well, and therefore we could infer the template strand position downstream nsp12. We found that the angle between the upstream template strand and the duplex product can only increase by ~33°, i.e. from ~87° to ~121°, before sterically clashing with nsp12 residues (**Fig. S10**). We suggest that the maximum angle is reached at ~30 pN, and further increasing the force does not further modify the template strand alignment, and therefore the nucleotide addition rate.

We show here that the polymerase incorporates nucleotides through three distinct catalytic pathways that we coined NAB, SNA and VSNA for nucleotide-additon burst, slow-nucleotide-addition (short pauses ≤ 1 s duration) and very-slow-nucleotide-addition (pauses ~1-5 s duration) catalytic pathways. The nucleotide addition cycle is conserved for all pathways and all starts with a thermally activated forward translocation, similarly to what has been observed with cellular DNA-dependent RNA polymerases, such as *Escherichia coli* RNA polymerase and the yeast RNA polymerase II (*27–29*). Following translocation, the nucleotide addition cycles carry on with nucleotide binding, and two successive irreversible steps, where the first is linked to the chemistry of nucleotide addition and the second to a large conformational change in the elongating polymerase-primer-template complex (**Fig. 6A**, **Fig. S9**). The force dependence of the second irreversible step kinetics indicates a global conformational change that displaces the RNA template between its exit and entry points into the polymerase complex by a distance equivalent to three bases (**Fig. 4**). What could this large conformational change be? Structural work on the related poliovirus and EV71 RdRps suggests that reopening of the active site at the end of the nucleotide addition cycle is concomitant with a large conformational change of the motif B loop and a displacement of the D-motif (*6, 45, 47*), with a rearrangement of the interactions between the template strand and the G-motif (*48*). Altogether, these rearrangements would release the “grip” of the polymerase on the primer-template to reset the polymerase and enable translocation at the next nucleotide addition cycle.

The structural origin of the SNA and VSNA pathways is difficult to assertain, as these pathways have not been previously identified. Mutating the G-motif to alter the interactions with the +1 position of the template strand significantly affects the kinetics of nucleotide addition (*48*). Because Pause 1 and Pause 2 probability increases with tension (**Fig. 4F**, **Fig. S5K**), one could hypothesize that destabilizing the aforementioned interaction lead the polymerase to the SNA/VSNA state. However, our previous study on poliovirus showed the error-prone mutant H273R of poliovirus RdRp induced a large increase in the VSNA pathway probability (*19*), though the mutation is distant from the catalytic site (*49*). A network of interactions between the polymerase motifs, co-factors and the primer-template likely lead the polymerase into the SNA and VSNA states in a stochastic manner. This states may be evolutionary conserved to regulate replication and transcription kinetics, but also to enable the surveillance of the coronavirus polymerase by other viral co-factors.

How the proofreading exonuclease, nsp14, detects mismatch incorporation by nsp12 RdRp is unknown. Is there a kinetic competition between nsp12 and nsp14? Does a pause in the elongating nsp12 provides the time needed for the exonuclease to intervene, as for some high-fidelity DNA polymerases (*22, 25*)? We have identified three pauses in the SARS CoV-2 RdRp elongation kinetics. While the long-lived backtrack pauses are catalytically incompetent, Pause 1 (SNA) and Pause 2 (VSNA) are the signatures of slow nucleotide incorporations and therefore represent good candidates fitting the kinetic fingerprint of nucleotide misincorporation. Pause 1 probability is too high to be consistent with mismatch incorporation, but Pause 2 kinetics are on par with pre-steady-state kinetic analysis of SARS-CoV-1 polymerase, where the average synthesis rate of a 10 nt product including a UTP:G mismatch was measured at (0.14 ± 0.01) s^−1^) in the presence of 50 μM NTP (*10*). At 25 pN and 50 μM NTP, we measured a similar exit rate for Pause 2 at (0.23 ± 0.01) s^−1^) (**Table S1**). Pause 2 probability at saturating concentration of NTP is similar to what we measured for poliovirus RdRp, where Pause 2 was identified as the kinetic signature of nucleotide mismatch incorporation (*19*). This would suggest that the fidelity of the coronavirus polymerase is on par with other viral RdRps. Furthermore, the life time of Pause 2, 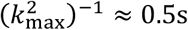 indicates the lower bound time interval for nsp14 to intervene. Future studies investigating the elongation kinetics of a – yet to identify – low fidelity mutant of the SARS-CoV-2 polymerase will clearly identify the pathway to mismatch incorporation and whether it is associated with the VSNA pathway.

We have significantly improved the stability of our magnetic tweezers assay for the noise not to be limited by the drift for an observation window as large as ~100 s, i.e. a ~50-fold improvement compared to the best resolution previously reported (*40–42*) (**Fig. S7**). We leveraged the better stability of our assay to directly monitor SARS-CoV-2 polymerase backtracking as deep as ~30 nt upstream the last incorporated nucleotide (**Fig. 5CD**, **Fig. S8**). Our result supports a recent cryo-EM study on a pre-assembled backtracked coronavirus RdRp complex (*43*). The dwell-time distribution of pauses resulting from polymerase backtrack is mathematically described by a power law with a −3/2 exponent (*34*), which is consistent with our findings for the long-lived third pause. Altogether, our direct observation and dwell time analysis support polymerase backtracking as origin for the long-lived pauses, which further generalizes backtracking as a common property in the viral RdRp world (*19, 30, 31*). We show here that SARS-CoV-2 polymerase backtracking is strongly stimulated by the presence of secondary structure downstream the polymerase (**Fig. 5F**). Given how heavily structured the coronavirus genome is (*39*), polymerase backtracking would likely be ubiquitous during coronavirus genome replication and transcription without the assistance of other viral co-factors. The helicase nsp13 or the single-stranded RNA binding protein nsp9 are good candidates for this assistive function, and several cryo-EM studies have reported that nsp9 and nsp13 associate to the coronavirus polymerase (*46, 50, 51*). A very recent study even identifies nsp13 as an essential component to establish the polymerase into a stable backtrack state (*43*). Therefore, nsp13 potentially modulates coronavirus polymerase backtracking, which may be essential to promote strand switching during viral genome recombination and transcription (*52, 53*).

Having characterized the three catalytic pathways of the coronavirus polymerase, it is now possible to understand by which pathway it incorporates nucleotide analogues. Of most importance, we show that the entry into any of the slow nucleotide addition pathways occurs before nucleotide (analogue) binding, and therefore reaction conditions (as NTP concentration) are extremely important to ensure proper evaluation of how nucleotide analogues are incorporated and the yield of incorporation. In the companion study whe leverage the bench-mark provided here to fingerprint the action of various nucleotide analogues, demonstrating why Remdesivir is better incorporated than T-1106 (a chemically more stable version of Favipiravir) (*9*). This result likely explains the higher efficacy of Remdesivir against SARS-CoV-2 infected cells than Favipiravir. Being able to tune the polymerase into a drug-incorporation competent pathway using small molecules binding may represent an avenue to further improve drug efficacy.

Replication and transcription is at the heart of coronavirus life-cycle, and form an important target for drug development. However, a mechanistic understanding of these processes has been lacking, forming an obstacle to efficiently target the key aspect that regulates viral genome synthesis in infected cells. Here, we provide a complete mechanistic framework describing the coronavirus core polymerase nucleotide addition cycle. We will leverage this single molecular platform to elucidate the structure-function-dynamics relationship of the multi-subunits RdRp complexes processing the coronavirus genome, offering new targets for drug development.

## Acknowledgements

DD was supported by the Interdisciplinary Center for Clinical Research (IZKF) at the University Hospital of the University of Erlangen-Nuremberg, the German Research Foundation grant DFG-DU-1872/3-1 and BaSyC – Building a Synthetic Cell” Gravitation grant (024.003.019) of the Netherlands Ministry of Education, Culture and Science (OCW) and the Netherlands Organisation for Scientific Research (NWO). DD thanks OICE for providing office, lab space, and access to their molecular biology lab. RNK was supported by grant AI123498 from NIAID, NIH. BC acknowledge grants by the Fondation pour la Recherche Médicale (Aide aux équipes), the SCORE project H2020 SC1-PHE-Coronavirus-2020 (grant#101003627), and the REACTing initiative (REsearch and ACTion targeting emerging infectious diseases). JJA and CEC were supported by grant AI045818 from NIAID, NIH.

## Author contribution

JJA, BC, CEC and DD conceived the research. SCB and MS performed the single molecule experiments. SCB, MS, MD and DD analyzed the data. PvN wrote the MLE analysis routine. FSP made the RNA used for the study. YW helped developing the magnetic tweezers autofocus. RNK provided SARS-CoV-2 polymerase proteins. MD performed theoretical analysis and modelling. SQ derived the structural model of the polymerase. All the authors have discussed and interpreted the results. All the authors have participated in writing the article.

