## Supplementary Materials for "The nucleotide addition cycle of the SARS-CoV-2 polymerase"

**This PDF file includes:**

Materials and Methods

Figures S1 to S10

Table S1

### Materials and Methods

#### *High-throughput magnetic tweezers apparatus*

The high-throughput magnetic tweezers used in this study have been described in detail elsewhere (*1*). Shortly, a pair of vertically aligned permanent magnets (5 mm cubes, SuperMagnete, Switzerland) separated by a 1 mm gap are positioned above a flow cell (see paragraph below) which is mounted on a custom-built inverted microscope. The vertical position and rotation of the magnets are controlled by two linear motors, M-126-PD1 and C-150 (Physik Instrumente PI, GmbH & Co. KG, Karlsruhe, Germany), respectively. The field of view is illuminated through the magnets gap by a collimated LED-light source, and is imaged onto a large chip CMOS camera (Dalsa Falcon2 FA-80-12M1H, Stemmer Imaging, Germany) using a 50× oil immersion objective (CFI Plan Achro 50 XH, NA 0.9, Nikon, Germany) and an achromatic doublet tube lens of 200 mm focal length and 50 mm diameter (Qioptic, Germany). To control the temperature, we used a system described in details in Ref. (*2*). A flexible resistive foil heater with an integrated 10 MΩ thermistor (HT10K, Thorlabs) is wrapped around the microscope objective and further insulated by several layers of kapton tape (KAP22-075, Thorlabs). The heating foil is connected to a PID temperature controller (TC200 PID controller, Thorlabs) to adjust the temperature within ~0.1 °C.

#### *Ultra-stable magnetic tweezers*

To improve the stability of the instrument, we introduced an autofocus (i.e. along the z-axis) to the magnetic tweezers assay. We hypothesized that reference bead (a surface melted polystyrene bead) subtraction alone could not fully correct for large drift (> 100 nm) during long timescale (> 20 minutes) measurements. Therefore, correction for the drift by establishing an autofocus with a high resolution piezo nanopositioner would correct for such detrimental effect. To achieve this, we selected a suitable surface-attached reference bead in the field of view and considered it for autofocus in the tracking algorithm. During the acquisition, the current z-axis position (average of last 60 z-positions) of the reference bead was compared every 1 sec interval against its target z-axis position (average of 60 z-positions at the beginning/suitable part of measurement). If the difference between the current and the target z-positions was larger than 0.5 nm, the drift/difference was subsequently corrected by adjusting the position of the objective using the piezo nanopositioner. Of note, the drift in our assay is much smaller than 0.5 nm/s. To quantitatively evaluate the improvement introduced by the autofocus into the measurement, we extracted the Allan deviation (AD) of a reference bead either with or without autofocus correction and either drift-corrected or not, i.e. by further subtracting the position of another reference bead. We estimated the AD at two different acquisition frequency, i.e. 58 and 500 Hz, using 100x microscope objective magnification (60 nm pixel size in the image plane) (**Fig. S7A-C**). The AD compares the noise in the bead position for partially overlapping, successive time windows of increasing duration τ. Therefore, the AD quantifies the magnitude of the noise when average over a given time interval, and informs on its origin (*3, 4*). For shot-noise limited measurements, the AD decreases as $1/\sqrt{\tau}$, to eventually increases when other sources of noise, e.g. mechanical drift, cumulates and dominates at larger τ. Therefore, the minimum of AD indicates the time interval over which the assay is stable and the best resolution achievable. This is clearly visible in **Fig. S7D**, where the AD at 58 Hz acquisition frequency reaches a minimum of AD $\sim 0.1\ nm$ in absence of reference bead subtraction at $\tau \sim 1\ s$. This minimum decreases to AD $\sim 0.01\ nm$ at $\tau \sim 30\ s$ when subtracting the reference bead position. Applying the autofocus, the drift is further reduced, with a minimum of AD$\sim 0.008\ nm$ at $\tau \sim 100\ s$, indicating a significant improvement in stability. Similar results were obtained at 500 Hz (**Fig. S7E**). We note that the resolution does not significantly improve below AD$\sim 0.008\ nm$, as we likely reached a physical limit in what is achievable in position correction for our assay.

*Recombinant Protein Expression of RdRp (nsp12) and cofactors (nsp7 and nsp8) from SARS-CoV-2*

This protocol was described in Ref. (*5*). SARS-CoV-2 nsp12: The SARS-CoV-2 nsp12 gene was codon optimized and cloned into pFastBac with C-terminal additions of a TEV site and strep tag (Genscript). The pFastBac plasmid and DH10Bac E. coli (Life Technologies) were used to create recombinant bacmids. The bacmid was transfected into Sf9 cells (Expression Systems) with Cellfectin II (Life Technologies) to generate recombinant baculovirus. The baculovirus was amplified through two passages in Sf9 cells, and then used to infect 1 L of Sf21 cells (Expression Systems) and incubated for 48 hrs at 27°C. Cells were harvested by centrifugation, resuspended in wash buffer (25 mM HEPES pH 7.4, 300 mM NaCl, 1 mM $MgCl_2$, 5 mM DTT) with 143 μl of BioLock per liter of culture. Cells were lysed via microfluidization (Microfluidics). Lysates were cleared by centrifugation and filtration. The protein was purified using StrepTactin Superflow agarose (IBA). StrepTactin eluted protein was further purified by size exclusion chromatography using a Superdex 200 Increase 10/300 column (GE Life Sciences) in 25 mM HEPES, 300 mM NaCl, 100 μM $MgCl_2$, 2 mM TCEP, at pH 7.4. Pure protein was concentrated by ultrafiltration prior to flash freezing in liquid nitrogen. SARS-CoV-2 nsp7 and nsp8: The SARS-CoV-2 nsp7 and nsp8 genes were codon optimized and cloned into pET46 (Novagen) with an N-terminal 6x histidine tag, an enterokinase site, and a TEV protease site. Rosetta2 pLys E. coli cells (Novagen) were used for bacterial expression. Cultures were grown to an OD600 of 0.8 and induced with a final concentration of 0.5 mM isopropyl β-D-1-thiogalactopyranoside (IPTG) and growth temperature was reduced to 16°C for 16 hrs. Cells were harvested by centrifugation and pellets were resuspended in wash buffer (10 mM Tris pH 8.0, 300 mM NaCl, 30 mM imidazole, 2 mM DTT). Cells were lysed via microfluidization and lysates were cleared by centrifugation and filtration. Proteins were purified using Ni-NTA agarose beads (Qiagen) and eluted with wash buffer containing 300 mM imidazole. Eluted proteins were digested with 1% w/w TEV protease during overnight room temperature dialysis (10 mM Tris pH 8.0, 300 mM NaCl, 2 mM DTT). Digested proteins were passed back over Ni-NTA to remove undigested protein before concentrating the proteins by ultrafiltration. Nsp7 and nsp8 proteins were further purified by size exclusion chromatography using a Superdex 200 Increase 10/300 column (GE Life Sciences). Purified proteins were concentrated by ultrafiltration prior to flash freezing with liquid nitrogen.

##### *Construct fabrication.*

The fabrication of the RNA hairpin has been described in detail in Ref. (*6*). The RNA hairpin is made of a 499 bp double-stranded RNA stem terminated by a 20 nt loop that is assembled from three ssRNA annealed together (**Figure S1A**), and two handles, one of 856 bp at the 5' end and one 822 bp at the 3' end. The handles include either a 343 nt digoxygenin-labeled ssRNA or a 443 nt biotin-labeled ssRNA (**Figure S1A**). Upon applied force above ~21 pN, the hairpin opens and frees a 1043 nt ssRNA template for SARS-CoV-2 polymerase (**Figure S1B**). To obtain the different parts of the RNA construct, template DNA fragments were amplified via PCR, purified (Monarch PCR and DNA cleanup kit) and in vitro transcribed (NEB HiScribe™ T7 High Yield RNA Synthesis Kit). Transcripts were then treated with Antarctic Phosphatase and T4 Polynucleotide Kinase. RNAs were purified using the RNA Clean & Concentrator-25 kit (Zymo Research). Individual RNA fragments were annealed and ligated with T4 RNA ligase 2 (NEB) to assemble the RNA hairpin.

RNA template for SARS-CoV-2 magnetic tweezers experiments (highlighted in yellow and in grey are the loop and the ssRNA template preceding the hairpin stem where the polymerase loads, respectively)

GUUCUACAUAGCGUGCAGACGUGAAUUUAAUCUCGCUGACGUGUAGACACAGUGCGUCU
GCUGUCGGGUCCCUCUGGUGACUGGGUAGUUGGACUUGCCCUUGGAAGACAUAGCAAGA
CCCUGCCUCUCUAUUGAUGUCACGGCGAAUGUCGGGGAGACAGCAGCGGCUGCAGACAUC
AGAUCGGAGUAAUACUCUCCGUAACUGGCCUUCUCUGAAUUCCGACGUUGUUAAGAUGG
CAGAGCCCGGUAAUCGCUACUUGACCAGAUAAGCUUUCCGUGGAUGGUUUAGAGGAAUC
ACAUCCAAGACUGGCUAAGCACGAAGCAACUCUUGAGUGUAAAAUUGUUGUCUCCUGUA
UUCGGGAUGCGGGUACUAGAUGACUGCAGGGACUCCGACGUUAAGUACAUUACCCCGUCA
UAGGCGCCGUUCAGGAUCACGUUACCGCCAUAAGAUGGGAGCAUGACUUCUUCUCCGCUG
CGCCCACGGAUCCAGUAGUGAUUAACAUUCGACAGCAUGCGCACUAAUCACUACUGGAUC

CGUGGGCGCAGCGGAGAAGAAGUCAUGCUCCCAUCUUAUGGCGGUAACGUGAUCCUGAAC
GGCGCCUAUGACGGGGUAAUGUACUUAACGUCGGAGUCCCUGCAGUCAUCUAGUACCCGC
AUCCCGAAUACAGGAGACAACAAUUUUACACUCAAGAGUUGCUUCGUGCUUAGCCAGUCU
UGGAUGUGAUUCCUCUAAACCAUCCACGGAAAGCUUAUCUGGUCAAGUAGCGAUUACCG
GGCUCUGCCAUCUUAACAACGUCGGAAUUCAGAGAAGGCCAGUUACGGAGAGUAUUACU
CCGAUCUGAUGUCUGCAGCCGCUGCUGUCUCCCCGACAUUCGCCGUGACAUCAAUAGAGA
GGCAGGGUCUUGCUAUGUCUUCCAAGGGCAAGUCCAACUACCCAGUCACCAGAGGGACCC
GACAGCAGACGCACUGUGUCUACACGUCAGCGAGAUUAAAUUCACGUCUGCACGCUAUGU
AGAACCCUCAGCCAACUCGGUCGCGUCGGA

The template contains 250 U (24%), 253 A (24%), 273 C (26%) and 267 G (26%).

#### *Flow-cell assembly*

The fabrication procedure for flow cells has been described in details in (*1*). To summarize, we sandwiched a double layer of Parafilm by two #1 coverslips, the top one having one hole at each end serving as inlet and outlet, the bottom one being coated with a 0.01% m/V nitrocellulose in amyl acetate solution. The flow cell is mounted into a custom-built holder and rinsed with ~1 ml of 1x phosphate buffered saline (PBS). 3 μm diameter polystyrene reference beads are attached to the bottom coverslip surface by incubating 100 μl of a 1:1000 dilution in PBS (LB30, Sigma Aldrich, stock conc.: 1.828*1011 particles per milliliter) for ~3 minutes. The tethering of the magnetic beads by the RNA hairpin construct relies on a digoxygenin/anti-digoxygenin and biotin-streptavidin attachment at the coverslip surface and the magnetic bead, respectively. Therefore, following a thorough rinsing of the flow cell with PBS, 50 μl of anti-digoxigenin (50 μg/ml in PBS) is incubated for 30 minutes. The flow cell was flushed with 1 ml of 10 mM Tris, 1 mM EDTA pH 8.0, 750 mM NaCl, 2 mM sodium azide buffer to remove excess of anti-digoxigenin followed by rinsing with another 0.5 ml of 1x TE buffer (10 mM Tris, 1 mM EDTA pH 8.0 supplemented with 150 mM NaCl, 2 mM sodium azide). The surface is then passivated by incubating bovine serum albumin (BSA, New England Biolabs, 10 mg/ml in PBS and 50% glycerol) for 30 minutes, and rinsed with 1x TE buffer.

#### *Single molecule SARS-CoV-2 polymerase activity experiments*

20 μl of streptavidin coated Dynal Dynabeads M-270 streptavidin coated magnetic beads (ThermoFisher) was mixed with ~0.1 ng of RNA hairpin (total volume 40 μl) and incubated for ~5 minutes before rinsing with ~2 ml of 1x TE buffer to remove any unbound RNA and the magnetic beads in excess. RNA tethers

were sorted for functional hairpins by looking for the characteristic jump in extension length due to the sudden opening of the hairpin during a force ramp experiment (*6*). The flow cell was subsequently rinsed with 0.5 ml reaction buffer (50 mM HEPES pH 7.9, 2 mM DTT, 2 µM EDTA, 5 mM $MgCl_2$). After starting the data acquisition at a force that would keep the hairpin open, 100 µl of reaction buffer containing the indicated concentrations of nsp12, nsp7, nsp8 (1:3:3 stoichiometry) and NTPs were flushed in the flow cell to start the reaction. For the pre-assembled polymerase experiments, 0.6 µM nsp12, 1.8 µM nsp7 and nsp8 were incubated for five minutes in the flow cell, while applying 35 pN force on the tether. The excess polymerase proteins were subsequently flushed away with 0.3 ml of reaction buffer (flow cell volume ~ 40 µl), followed by the injection of 100 µl of reaction buffer with 500 µM NTP. The experiments were conducted at a constant force as indicated for a duration of 20 to 60 minutes. The camera frame rate was fixed at 58 Hz and the temperature set to 25°C. A custom written Labview routine controlled the data acquisition and the (x-, y-, z-) positions analysis/tracking of both the magnetic and reference beads in real-time (*7*). Mechanical drift correction was performed by subtracting the reference bead position from the magnetic bead positions.

#### *Data processing*

The activity traces of SARS-CoV-2 polymerase converts the tether from ssRNA to dsRNA, which concomitantly decreases the end-to-end extension of the tether. The change in extension measured in micron was subsequently converted into replicated nucleotides $N_R$, low-pass filtered at 2 Hz and the dwell times were extracted using a dwell time window of 10 nt as described in Ref. (*8*). For data acquired on the closed hairpin, the data was converted into nucleotides based on the average increase in extension for completed activity traces and low-pass filtered at 1 Hz. Dwell times were extracted as for the data on the ssRNA. The dwell times of all the traces for a given experimental condition were assembled and further analyzed using a maximum likelihood estimation (MLE) fitting routine to extract the parameters from the stochastic-pausing model (see below).

#### *SARS-CoV-2 activity trace product length analysis*

To extract the product length of the polymerase complex, only the traces where the beginning and the end could clearly be distinguished and for which the tether did not rupture for ten minutes following the last observed elongation activity were considered. We represented the mean product length, as well as one standard deviation of the mean from 1000 bootstraps as error bars.

#### *Model fitting*

There are many kinetic models that are consistent with the empirical dwell-time distributions we observe. The first thing we need to account for is multiple entry into a pause state in one dwell-time window.

##### *Stochastic pausing model*

For concreteness, we first consider the situation where we have three characteristic times (corresponding to nucleotide addition, and two to two different pauses) in a single nucleotide addition step. Based on the data, we assume that we have separation of timescales $\tau_0 \ll \tau_1 \ll \tau_2$, and that each process (nucleotide addition, or pausing) dominates the single-nt dwell-time distribution for times around its characteristic timescale. This assumption washes out most details of the kinetic scheme that connects the pauses with nucleotide addition, but allows us to determine the general form of the dwell-time distribution without specifying how the pauses are connected to the nucleotide addition pathway. Under the above assumptions, we have the approximate single-nt dwell-time distribution

$$P_{1\mathrm{nt}}(t) = \frac{p_0}{\tau_0} e^{-\frac{t}{\tau_0}} + \frac{p_1}{\tau_1} e^{-\frac{t}{\tau_1}} + \frac{p_2}{\tau_2} e^{-\frac{t}{\tau_2}}, \qquad \tau_0 < \tau_1 < \tau_2, \qquad p_0 + p_1 + p_2 = 1.$$

Moving over to Laplace space this becomes

$$\psi_{1\mathrm{nt}}(s) = \int_0^\infty \mathrm{d}t\, P_{1\mathrm{nt}}(t) e^{-st} = \frac{p_0}{1+\tau_0 s} + \frac{p_1}{1+\tau_1 s} + \frac{p_2}{1+\tau_2 s},$$

Using the fact that convolutions in real space are products in Laplace space, the first passage time distribution across a $N$ nucleotide window can be written as

$$\psi_{N\mathrm{nt}} = \left(\frac{p_0}{1+\tau_0 s} + \frac{p_1}{1+\tau_1 s} + \frac{p_2}{1+\tau_2 s}\right)^N$$

$$= \sum_{n_0=0}^{N} \sum_{n_1=0}^{N-n_0} \binom{N}{n_0, n_1, N-n_0-n_1} p_0^{n_0} p_1^{n_1} p_2^{N-n_0-n_1} \left(\frac{1}{1+\tau_0 s}\right)^{n_0} \left(\frac{1}{1+\tau_1 s}\right)^{n_1} \left(\frac{1}{1+\tau_2 s}\right)^{N-n_0-n_1}$$

$$= \underbrace{p_0^N \left(\frac{1}{1+\tau_0 s}\right)^N}_{\text{only fast timescale}} + \underbrace{\sum_{n_0=0}^{N-1} \binom{N}{n_0, N-n_0, 0} p_0^{n_0} p_1^{N-n_0} \left(\frac{1}{1+\tau_0 s}\right)^{n_0} \left(\frac{1}{1+\tau_1 s}\right)^{N-n_0}}_{\text{at least one intermediate but no slow timescale}}$$

$$+ \underbrace{\sum_{n_0=0}^{N-1} \sum_{n_1=0}^{N-n_0-1} \binom{N}{n_0, n_1, N-n_0-n_1} p_0^{n_0} p_1^{n_1} p_2^{N-n_0-n_1} \left(\frac{1}{1+\tau_0 s}\right)^{n_0} \left(\frac{1}{1+\tau_1 s}\right)^{n_1} \left(\frac{1}{1+\tau_2 s}\right)^{N-n_0-n_1}}_{\text{at least one slow timescale}}$$

In the last step we have separated out the terms that contain only the fast process (first term), at least one of the intermediate process but none of the slow process (second term), and those that contain at least one slow process (third term).

This means that we do not have to keep track of the fast timescale when considering the second term, nor the fast and intermediate timescale when considering the third term. We approximate

$$\psi_{N\mathrm{nt}} \approx \underbrace{p_0^N \left(\frac{1}{1+\tau_0 s}\right)^N}_{\psi_{N\mathrm{nt}}^0} + \underbrace{\sum_{n_0=0}^{N-1} \binom{N}{n_0, N-n_0} p_0^{n_0} p_1^{N-n_0} \left(\frac{1}{1+\tau_1 s}\right)^{N-n_0}}_{\psi_{N\mathrm{nt}}^1}$$

$$+ \underbrace{\sum_{n_0=0}^{N-1} \sum_{n_1=0}^{N-n_0-1} \binom{N}{n_0, n_1, N-n_0-n_1} p_0^{n_0} p_1^{n_1} p_2^{N-n_0-n_1} \left(\frac{1}{1+\tau_2 s}\right)^{N-n_0-n_1}}_{\psi_{N\mathrm{nt}}^2}$$

This approximation breaks down in the short time limit ($t \ll \tau_0$) as the two last terms no not account for that nucleotide addition is needed to traverse the window even when there are pauses in the dwell-time window (see below).

The experimental dwell time distributions are all well fit with exponential shoulders for Pause 1 and Pause 2 using a maximum-likelihood approach. This corresponds to exchanging the second ($\psi_{N\mathrm{nt}}^1$) and third ($\psi_{N\mathrm{nt}}^2$) term in the above with simple exponential processes that capture the probability and the average time of each pause. The probability and average time can respectively be written as

$$q_i = \int_0^\infty \mathrm{d}t\, P_{N\mathrm{nt}}(t) = \psi_{N\mathrm{nt}}^i(0), \qquad T_i = \frac{\int_0^\infty \mathrm{d}t\, t\, P_{1\mathrm{nt}}(t)}{\int_0^\infty \mathrm{d}t\, P_{1\mathrm{nt}}(t)} = -\partial_s \ln \psi_{N\mathrm{nt}}^i(0), \qquad i = 1, 2.$$

The estimated weights and timescales over the $N$nt can be calculated as

$$q_1 = (p_0 + p_1)^N - p_0^N, \qquad T_1 = \frac{N p_1 (p_0 + p_1)^{N-1}}{(p_0 + p_1)^N - p_0^N} \tau_1$$

$$q_2 = 1 - (p_0 + p_1)^N, \qquad T_2 = \frac{N p_2}{1 - (p_0 + p_1)^N} \tau_2.$$

Based on this, the fitting function would be

$$P_{N\mathrm{nt}}(t) \approx \frac{1 - q_1 - q_2}{\tau_0 (N-1)!} \left({}^{t}/_{\tau_0}\right)^{N-1} e^{-t/\tau_0} + Q(t) \left(\frac{q_1}{T_1} e^{-t/T_1} + \frac{q_2}{T_2} e^{-t/T_2}\right)$$

where we have introduced the regularizing function

$$Q(t) = \frac{(t/(\tau_0 N_{\mathrm{dw}}))^{N_{\mathrm{dw}}-1}}{1 + (t/(\tau_0 N_{\mathrm{dw}}))^{N_{\mathrm{dw}}-1}}$$

to account for the fact that even when there is a pause, the short timescales are still dominated by a rapid succession on nucleotide addition steps. The fit results dependence on these cut-offs is negligible as long as they are introduced in regions where the corresponding term is sub-dominant. Here the cut is placed under the center of the elongation peak, guaranteeing that it is placed where pausing is sub-dominant.

We can now translate between the probabilities and timescales over a $N$nt window to the 1 nt window through

$$p_0 = \left(1 - (q_1 + q_2)\right)^{1/N}, \qquad p_1 = (1 - q_2)^{1/N} - \left(1 - (q_1 + q_2)\right)^{1/N}, \qquad p_2 = 1 - (1 - q_2)^{1/N}$$

$$\tau_1 = \frac{(p_0 + p_1)^N - p_0^N}{N p_1 (p_0 + p_1)^{N-1}} T_1, \qquad \tau_2 = \frac{1 - (p_0 + p_1)^N}{N p_2} T_2$$

Here, we explicitly also allow for a third, very improbable (never entered twice in a dwell-time window), and fat-tailed pause

$$P_{N\mathrm{nt}}(t) \approx \frac{1 - q_1 - q_2}{\tau_0 (N-1)!} \left({}^{t}/_{\tau_0}\right)^{N-1} e^{-t/\tau_0} + Q(t)\left(\frac{q_1}{T_1} e^{-t/T_1} + \frac{q_2}{T_2} e^{-t/T_2} + \frac{a_{\mathrm{bt}}}{2(1 + t/1s)^{3/2}}\right). \qquad \textbf{(S1)}$$

The additional third pausing term captures the asymptotic power-law decay (amplitude $a_{\mathrm{bt}}$) of the probability of dwell-times dominated by a backtrack. The backtracked asymptotic term needs to be further regularized for times shorter than the diffusive backtrack step. We have introduced a regularization at 1s, but the precise timescale does not matter, as long as it is set within the region where the exponential pauses dominate over the backtrack.

##### *Maximum likelihood estimation fitting routine*

The above stochastic-pausing model was fit to the dwell time distributions using a custom Python 3.7 routine. The dwell-time distribution is fit to the experimentally collected dwell-times $\{t_i\}_i$ by minimizing the likelihood function (*9*)

$$L = -\sum_i \ln P_{N\mathrm{nt}}(t_i) \qquad \textbf{(S2)}$$

with respect to timescales and probabilistic weights. We calculated the statistical error on the parameters by applying the MLE fitting procedure on 100 bootstraps of the original data set (*10*), and reported the standard deviation for each fitting parameters.

#### *Modelling of Pause 1 propensity under force and concentration sweeps*

In the main text we show that Pause 1 and Pause 2 are described by Michaelis-Menten kinetics (**Figure 4D**), and their probabilities increase with force (**Figure 3F**). We therefore hypothesize that they originate from the pre-translocated state of the polymerase. Pause 1 (p1) dominates in probability, and though we do not know how Pause 2 or other the long-lived pauses are connected to Pause 1, we can ignore their effect on the analysis of Pause 1 and elongation.

##### *The reaction kinetic scheme*

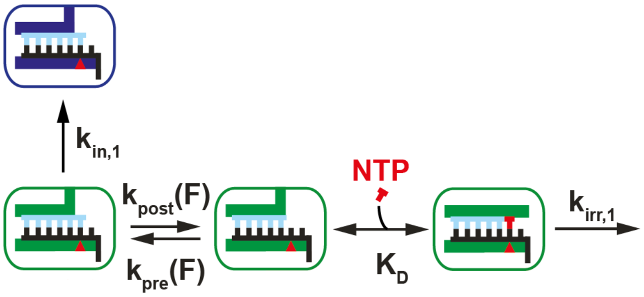

We consider here the working-model reaction scheme illustrated to the left. The nucleotide addition reaction starts with the polymerase fluctuating in position between the pre- and post-translocated states (rates $k_{\mathrm{post}}(F)$ and $k_{\mathrm{pre}}(F)$), and is stabilized forward upon nucleotide binding. Translocation is not assumed equilibrated, but binding is assumed equilibrated with dissociation constant $K_{\mathrm{D}}$. Once bound, the nucleotide is irreversibly incorporated at rate $k_{\mathrm{irr,1}}$.

We assume that all force-dependent rates can be written in terms of a shift of the transition state according to

$$k_{\mathrm{X}}(F) = k_{\mathrm{X}}(F_0)e^{-\delta_X(F-F_0)/k_{\mathrm{B}}T}$$

In the above, $\delta_X$ is the distance to the transition state in the direction of a growing transcript, $F$ is the tether tension and $F_0$ is an (arbitrary) reference tension.

By summing probabilities over paths, the first passage-time distribution of completing the irreversible step when starting from the pre-translocated state can be written in Laplace space as

$$\begin{aligned}
&\Psi_{\mathrm{irr,1}}(s) \\
&= \sum_{n=0}^{\infty}\left(\frac{k_{\mathrm{post}}(F)}{s + k_{\mathrm{in,1}} + k_{\mathrm{post}}(F)}\frac{K_{\mathrm{pre}}(F,[\mathrm{NTP}])}{s + K_{\mathrm{pre}}(F,[\mathrm{NTP}]) + K_{\mathrm{irr,1}}([\mathrm{NTP}])}\right)^n \\
&\times \frac{k_{\mathrm{post}}(F)}{s + k_{\mathrm{in,1}} + k_{\mathrm{post}}(F)}\frac{K_{\mathrm{irr,1}}([\mathrm{NTP}])}{s + K_{\mathrm{pre}}(F,[\mathrm{NTP}]) + K_{\mathrm{irr,1}}([\mathrm{NTP}])} \\
&= \frac{K_{\mathrm{irr,1}}([\mathrm{NTP}])k_{\mathrm{post}}(F)}{\left(s + k_{\mathrm{post}}(F) + k_{\mathrm{in,1}}\right)\left(s + K_{\mathrm{pre}}(F,\mathrm{NTP}]) + K_{\mathrm{irr,1}}([\mathrm{NTP}])\right) - k_{\mathrm{post}}(F)K_{\mathrm{pre}}(F,[\mathrm{NTP}])}.
\end{aligned}$$

In the equation above, we have defined the effective rates out of the binding equilibrated post-translocated state as

$$K_{\text{pre}}(F,[\text{NTP}]) = k_{\text{pre}}(F)\frac{K_{\text{D}}}{[\text{NTP}]+K_{\text{D}}}, \qquad K_{\text{irr},1}([\text{NTP}]) = k_{\text{irr},1}\frac{[\text{NTP}]}{[\text{NTP}]+K_{\text{D}}}.$$

For notational convenience we define the dimensionless rates

$$\gamma_{\text{in},1} = \frac{k_{\text{in},1}}{k_{\text{in},1}+k_{\text{post}}(F)}, \qquad \gamma_{\text{pre}} = \frac{k_{\text{pre}}(F)}{k_{\text{in},1}+k_{\text{post}}(F)}, \qquad \gamma_{\text{irr},1} = \frac{k_{\text{irr},1}}{k_{\text{in},1}+k_{\text{post}}(F)}.$$

*The observables*

From the first-passage time distribution we can deduce the probability to lock in the next base before entering the pause as

$$P_{\text{irr},1} = \Psi_{\text{irr},1}(0) = \left(1-\gamma_{\text{in},1}\right)\frac{[\text{NTP}]}{[\text{NTP}]+K_{\text{P}1}},$$

where we have introduced the modified dissociation constant

$$K_{\text{P}1} = K_{\text{D}}\frac{\gamma_{\text{pre}}\gamma_{\text{in},1}}{\gamma_{\text{irr},1}}$$

The inverse average time it takes to move from the pre-translocated state through the irreversible step can be written as

$$k_{\text{irr},1} = \left(-\frac{\partial \ln \Psi_{\text{irr},1}(s)}{\partial s}\bigg|_{s=0}\right)^{-1} = V_{\text{irr},1}^{\text{max}}\frac{[\text{NTP}]+K_{\text{P}1}}{[\text{NTP}]+K_{\text{D}}^{\text{app}}} \qquad \textbf{(S3)}$$

where

$$V_{\text{irr},1}^{\text{max}} = \frac{\gamma_{\text{irr},1}}{1+\gamma_{\text{irr},1}}, \qquad K_{\text{D}}^{\text{app}} = K_{\text{D}}\frac{1+\gamma_{\text{pre}}}{1+\gamma_{\text{irr},1}}.$$

*Catalytic rate*

We see no appreciable concentration dependence in the total catalytic rate at $F = 25$ pN (nucleotide addition rate, **Figure 3D**), which is only consistent with **Equation S3**) if $[\text{NTP}] \ll K_{\text{P}1}, K_{\text{D}}^{\text{app}}$. This implies that the probability to enter Pause 1,

$$P_{\text{P1}} = 1 - P_{\text{irr},1} = \frac{\gamma_{\text{in},1}[\text{NTP}] + K_{\text{P1}}}{[\text{NTP}] + K_{\text{P1}}}, \quad \textbf{(S4)}$$

is constant and close to unity, contradicting both the concentration dependence and quantitative values seen in **Figure 3D**. Thus, the irreversible step of our working-model reaction scheme cannot be rate-limiting, we conclude there is another irreversible, concentration independent, and rate-limiting step. This second, irreversible step (rate $k_{\text{irr},2}$), is associated with a large conformational change of the polymerase-primer-template complex.

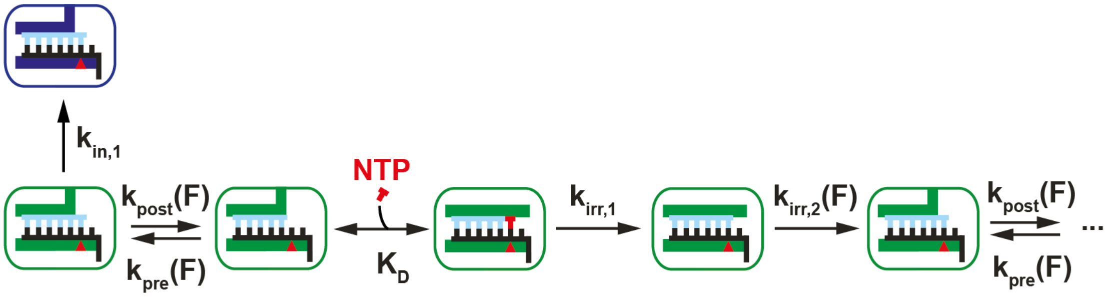

*Pause 1 probability*

Next, we turned to the pause probability. We performed a global fit of the pause probability $P_{\text{P1}}$ (**Equation S4**), over the concentration sweeps at tether tension $F = 25$ pN (**Figure 3E**, using all but lowest concentration point) and 35 pN (**Figure S5F**, using all but two lowest concertation points), and one force sweep at [NTP] = 0.5 mM (**Figure 3F**, using all but three highest force points). We minimized the error-weighted square deviation under the assumption that the distance to the transition state in the pre- to post-translocation $\delta > 0$, and that the distance from the other direction is $a_{\Delta\text{bp}} - \delta > 0$. From this we estimated (**Figure 6B-D**)

$$\delta = 0.0 \text{ nm}, \qquad a_{\Delta\text{bp}} = 0.23 \text{ nm}, \qquad \gamma_{\text{in},1} = 0.046, \qquad K_{\text{P1}} = 9.0 \ \mu\text{M}$$

at a tether tension of 25 pN. Allowing $\delta$ to be negative results in a very small negative value. In our data, we thus see no evidence that the pre- to post- translocation step is force dependent ($\delta \approx 0$ nm), while the post- to pre-translocation step acts over a distance $a_{\Delta\text{bp}} = \delta x_{\text{ss}} - \delta x_{\text{ds}}$ corresponding well to the difference in average length of a single- ($\delta x_{\text{ss}}$) and double-stranded ($\delta x_{\text{ds}}$) base in the present force range (*11*). We further see that translocation is not fast compared to NTP binding and pausing, as there is a non-zero probability ($\gamma_{\text{in},1} > 0$) to enter a pause even at saturating NTP concentrations; consequently, translocation is not equilibrated.

*The effect of a dsRNA structure downstream the polymerase*

In the presence of a hairpin, the energy of the post-translocated state is destabilized by the tension dependent melting energy

$$\epsilon(F) = \epsilon_0 - 2a_{\mathrm{ss}}F$$

of the outer hairpin base pair. Here $\epsilon_0$ is the relaxed melting energy and $a_{\mathrm{ss}}$ is the typical extension of a single-stranded (ss) base in our force range. Assuming that the distance to the transition state from pre- to post-translocated state is negligible also in the presence of a hairpin, only the rate from post- to pre-translocated state changes

$$k_{\mathrm{pre}}^{\mathrm{hp}}(F) = k_{\mathrm{pre}}^{\mathrm{no-hp}}(F)\exp(\epsilon(F)/k_{\mathrm{B}}T) \qquad \textbf{(S5)}$$

in the presence of hairpin. We have already fitted out $a_{\Delta\mathrm{bp}}$, the difference between a ss base and a ds base-pair in the present force range. dsRNA is $\sim$ 95% stretched in our force range, with a crystallographic length of 0.28 nm/bp (*12*), giving $\delta x_{\mathrm{ds}} \approx 27$ nm, and thus $a_{\mathrm{ss}} = 0.27\ \mathrm{nm} + a_{\Delta\mathrm{bp}} = 0.50$ nm; in accord with what is expected from the literature (*13*). Using this value, injecting **Equation S5** into **Equation S4**, we fitted Pause 1 probability in the presence of a hairpin (**Figure 6E**), and extract a zero-tension melting energy $\epsilon_0 = 18\ k_{\mathrm{B}}T$, i.e. a melting force of ~18 pN, which agrees well with the hairpin opening fully at ~22 pN (**Figure S1B**). We can also extract the probability to enter Pause 1 at zero force when the polymerase replicates through a dsRNA template, i.e. $P_{\mathrm{P1,hp}}(F = 0) \approx 0.31$; This should be compared to $P_{\mathrm{P1,no-hp}}(F = 0) \approx 0.049$ at the same conditions but without a hairpin.

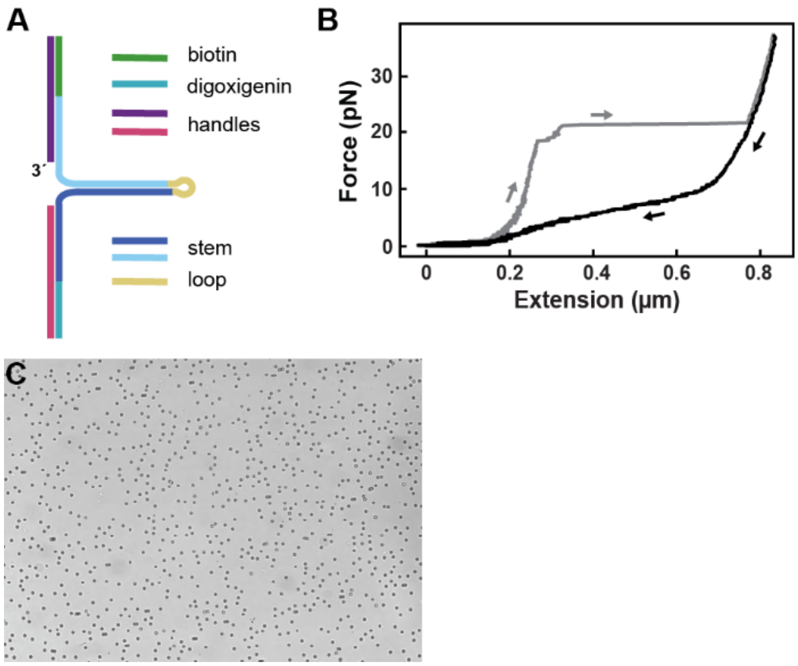

**Figure S1. Experimental conditions and data representation of SARS-CoV-2 high throughput magnetic tweezers experiments. (A)** Schematic of the RNA hairpin construct assembled from hybridizing and ligating single stranded RNAs. **(B)** Force as a function of the extension of the RNA hairpin presented in (A). Increasing (hairpin opening) and decreasing (hairpin closing) force ramp represented in grey and black, respectively. **(C)** Typical field of view containing ~450 hairpin tethered magnetic beads in a high-throughput magnetic tweezers assay.

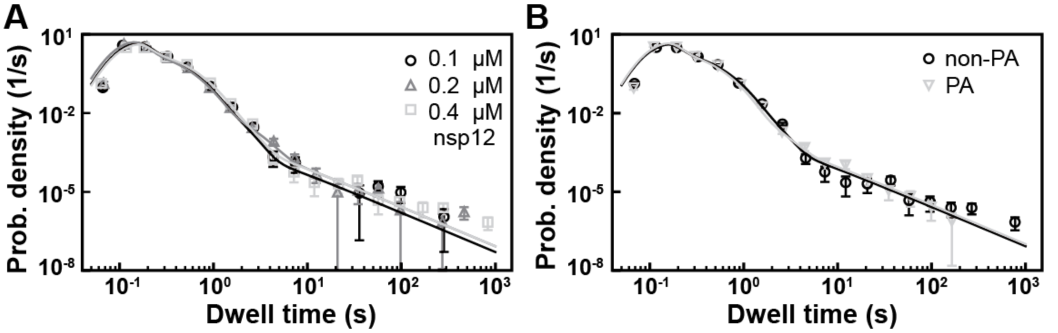

**Figure S2: SARS-CoV-2 elongation kinetics is not affected by different concentrations and pre-assembling of the polymerase.** All experiments were conducted at 35 pN and 25 °C. **(A)** Dwell time distributions of three different concentrations of nsp12 (0.1 µM (circle), 0.2 µM (triangle) and 0.4 µM (square)) and their corresponding MLE fits (solid lines). The stoichiometry of nsp12, nsp7, nsp8 is kept at 1:9:9. **(B)** The dwell time distributions of pre-assembled (PA) and non-pre-assembled (non-PA) SARS-CoV-2 polymerase. The error bars denote one standard deviation of 1000 bootstrap procedures.

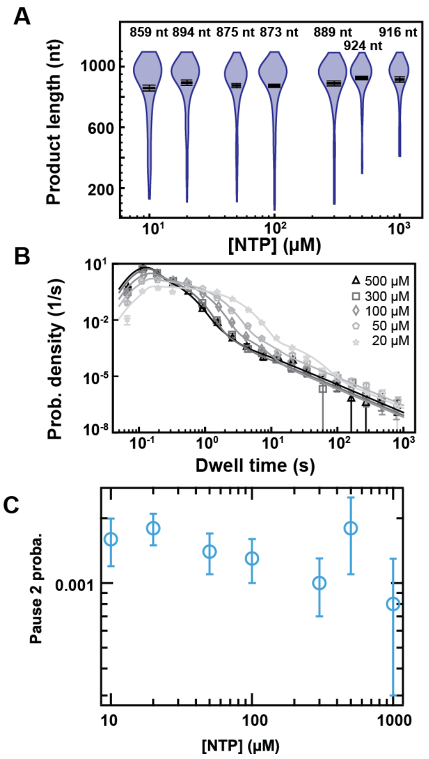

**Figure S3: SARS-CoV-2 elongation kinetics are affected by NTP concentration at 25 pN applied tension.** All data was acquired at 25 pN, 25 °C, 0.6 µM nsp12, 1.8 µM nsp7 and 1.8 µM nsp8. **(A)** The product length of SARS-CoV-2 polymerase complex at different NTP concentrations. The mean product lengths are denoted above the violin plots and represented as horizontal thick black lines, flanked by one standard deviation error bars extracted from 1000 bootstrap procedures. **(B)** The dwell time distributions at different NTP concentrations (500 µM (triangle), 300 µM (square), 100 µM (diamond), 50 µM (pentagon) and 20 µM (star)) and their corresponding MLE fits (solid lines). The error bars denote the standard deviation of 1000 bootstrap procedures. **(C)** Zoom-in of the Pause 2 probability as a function of NTP concentration and 25 pN applied force. The error bars denote the standard deviation of 100 bootstrap procedures.

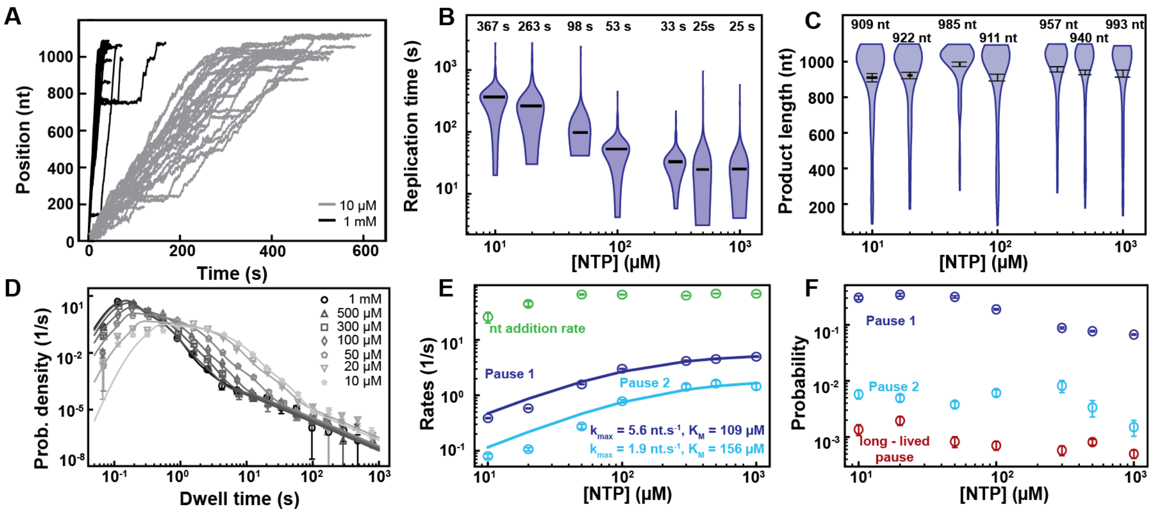

**Figure S4: Pause 1 and Pause 2 are the signature of slow catalytically competent states.** All experiments were conducted at 35 pN, 25 °C, 0.6 µM nsp12, 1.8 µM nsp7 and 1.8 µM nsp8. **(A)** Traces of SARS-CoV-2 polymerase complex activity at 1 mM (black) and 10 µM (grey) NTP. The **(B)** replication time and **(C)** product length of SARS-CoV-2 polymerase complex as a function of NTP concentration. The median replication times (B) and the mean product lengths (C) are denoted above the violin plots and represented as horizontal thick black lines, flanked by one standard deviation error bars extracted from 1000 bootstrap procedures. **(D)** The dwell time distributions at 1 mM (circles), 500 µM (triangles), 300 µM (squares), 100 µM (diamonds), 50 µM (pentagons), 20 µM (upside down triangle) and 10 µM (stars) and the corresponding MLE fits (solid lines). **(E)** Nucleotide addition rate (green), Pause 1 (dark blue) and Pause 2 (cyan) exit rates as a function of NTP concentration. The pause exit rates are fitted (solid lines) with the Michaelis-Menten equation. **(F)** The probabilities for Pause 1 (dark blue), Pause 2 (cyan) and backtrack (red). The error bars in (D) denote one standard deviation of 1000 bootstrap procedures. The error bars in (E) and (F) denote one standard deviation of 100 bootstrap procedures.

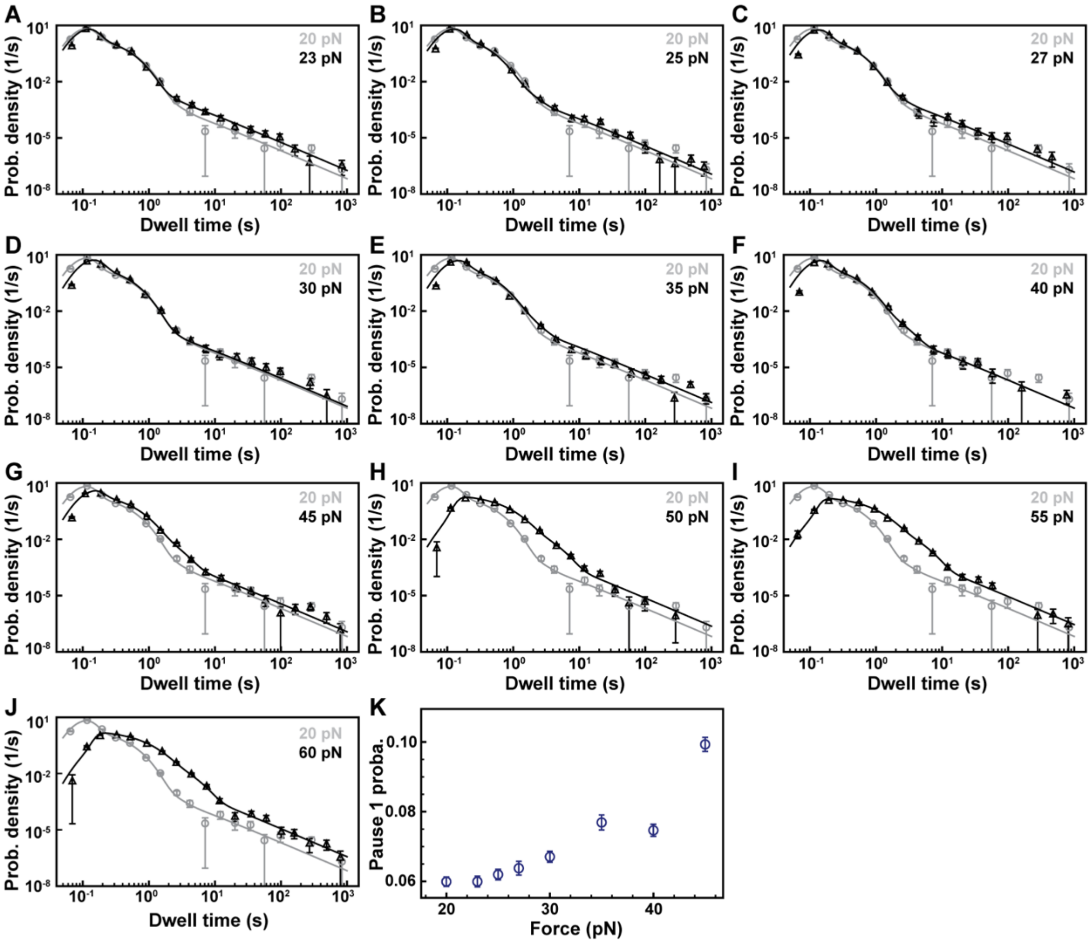

**Figure S5: Effects of applied tension on SARS-CoV-2 polymerase kinetics.** All experiments were conducted at 500 µM NTP, 25 °C, 0.6 µM nsp12, 1.8 µM nsp7 and 1.8 µM nsp8. The dwell time distributions of SARS-CoV-2 polymerase activity at different forces (**(A)** 23 pN, **(B)** 25 pN, **(C)** 27 pN, **(D)** 30 pN, **(E)** 35 pN, **(F)** 40 pN, **(G)** 45 pN, **(H)** 50 pN, **I)** 55 pN, **(J)** 60 pN) and the corresponding MLE fits (solid lines). The dwell time distribution at 20 pN (grey circles) and the corresponding MLE fit (solid line) is added to each panel to help guide the eye. **(K)** Pause 1 probability as a function of force. The error bars in (A-J) denote one standard deviation of 1000 bootstrap procedures and in (K) one standard deviation of 100 bootstrap procedures.

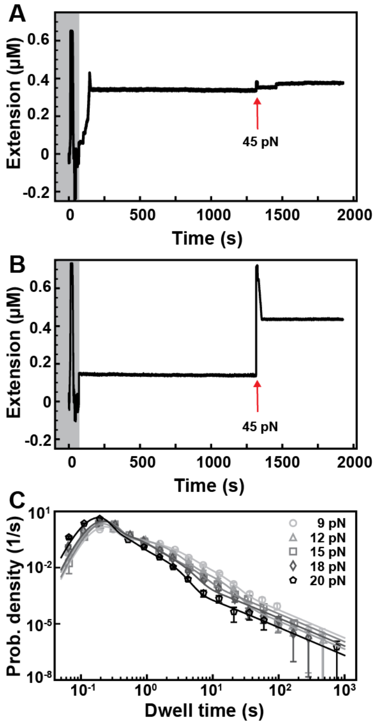

**Figure S6: SARS-CoV-2 polymerase kinetics on a dsRNA hairpin template.** All experiments were conducted at 500 μM NTP, 25 °C 0.2 μM nsp12, 1.8 μM nsp7 and 1.8 μM nsp8. Experiments were conducted such that first the hairpin was opened at a high force (45 pN) and then closed (4 pN) after which the nsps and 500 μM NTPs were flushed in. Just after flushing the magnet was moved to the measurement force. After 20 minutes, the force was increased to 45 pN to ensure the hairpin opening and immediately lowered to 30 pN (suitable force for polymerase activity). **(A)** A trace of SARS-CoV-2 polymerase activity at 18 pN on a closed hairpin. The increase in applied force to 45 pN shows only a small difference in end-to-end extension. **(B)** If no activity of the SARS-CoV-2 polymerase could be observed, an increase in applied force to 45 pN shows a jump in end-to-end extension due to the hairpin opening. Afterwards activity of the SARS-CoV-2 polymerase at 45 pN on the ssRNA can be observed. **(C)** The dwell time distributions at different applied tension 9 pN (circle), 12 pN (triangle), 15 pN (square), 18 pN (diamond) and 20 pN (pentagon)) and their corresponding MLE fits (solid lines). The error bars denote the standard deviation of 1000 bootstrap procedures.

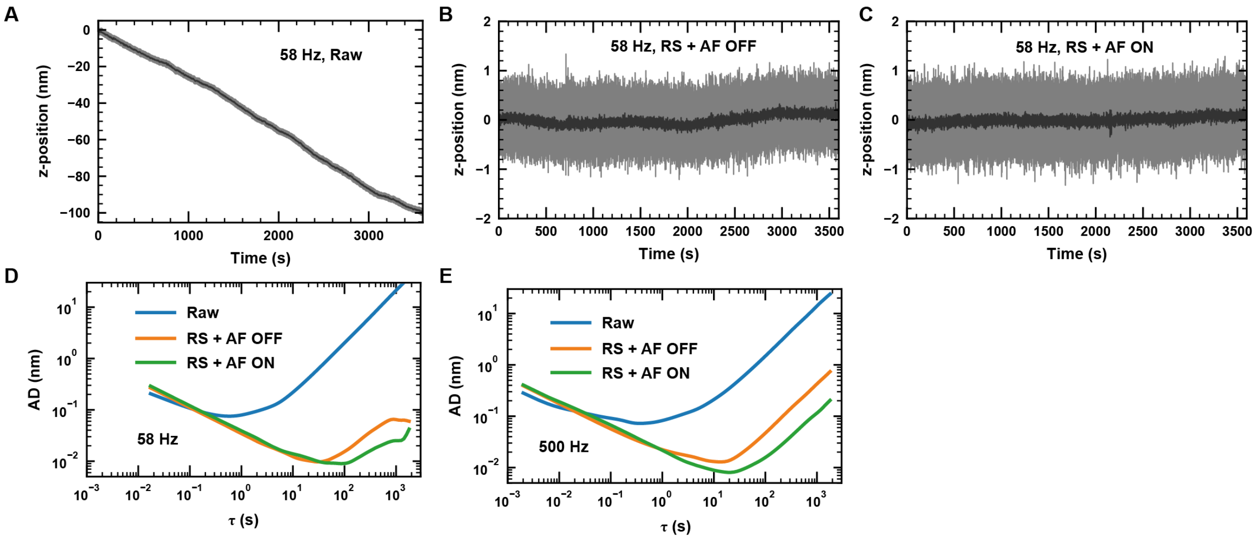

**Figure S7: Ultra-stable magnetic tweezers.** All the data were acquired using a 100x objective magnification and using 3 μm diameter surface-attached polystyrene reference beads. RS: reference subtracted; AF: auto-focus. **(A)** Raw data acquired at 58 Hz camera acquisition frequency (grey), low-pass filtered at 1 Hz (dark grey). **(B)** Same data, drift corrected by subtracting the z-position of another reference bead. **(C)** Data acquired while using the autofocus and drift corrected by subtracting the z-position of another reference bead. **(D)** Allan deviation (AD) of the traces in (A) (blue), (B) (orange) and (C) (green). **(E)** Same as in (D), now with data acquired at 500 Hz camera acquisition frequency.

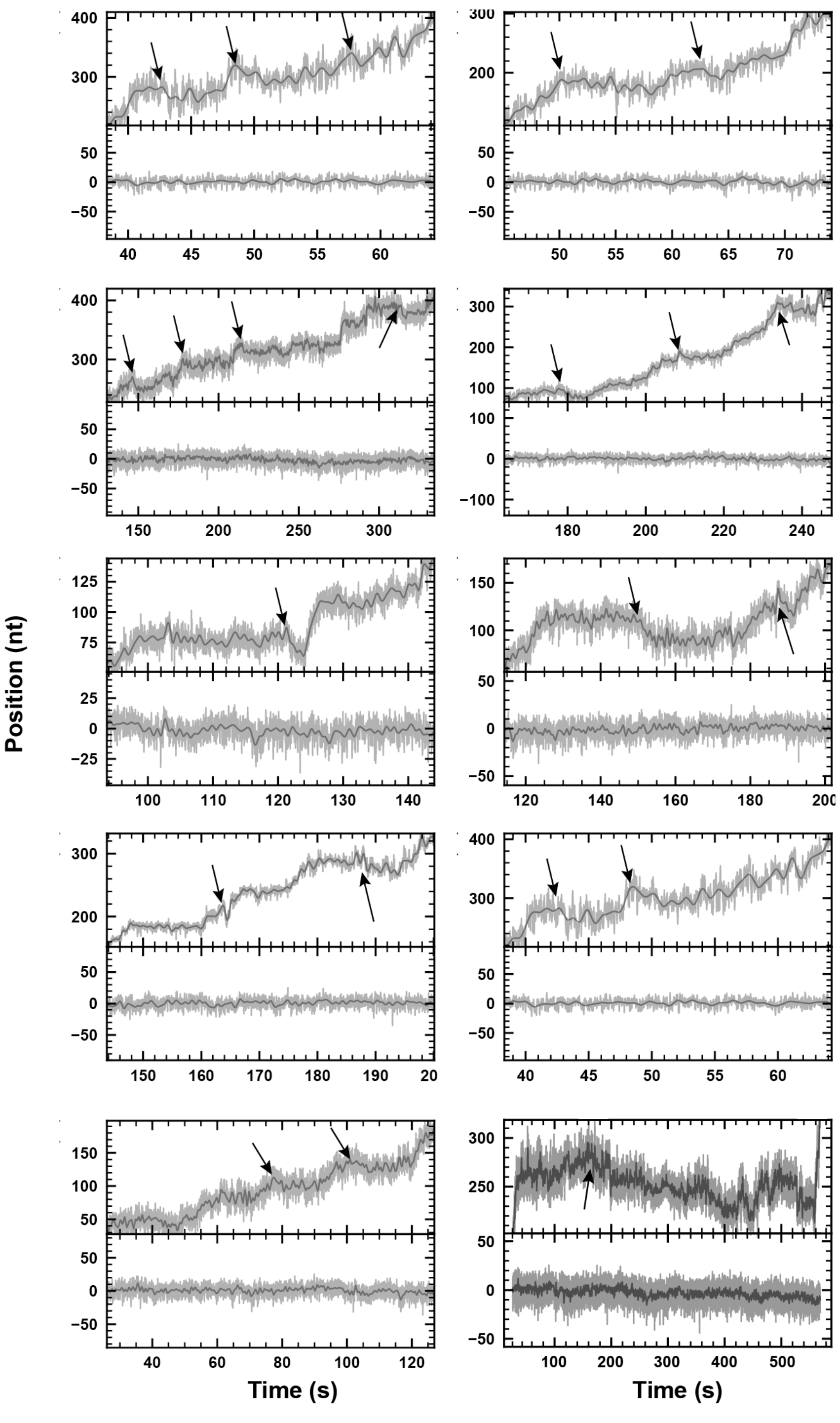

**Figure S8: Examples of SARS-CoV-2 polymerase backtrack.** Traces were acquired in 100x objective magnification and 9 pN force, using the autofocus as described in Supplementary Materials, and at 58 Hz camera acquisition frequency (grey), which was low-pass filtered at 1 Hz (dark grey). Each panel is divided in two parts: top, zoom in SARS-CoV-2 polymerase activity with black arrows indicating where the backtrack starts; bottom, trace of a tether without activity acquired simultaneously.

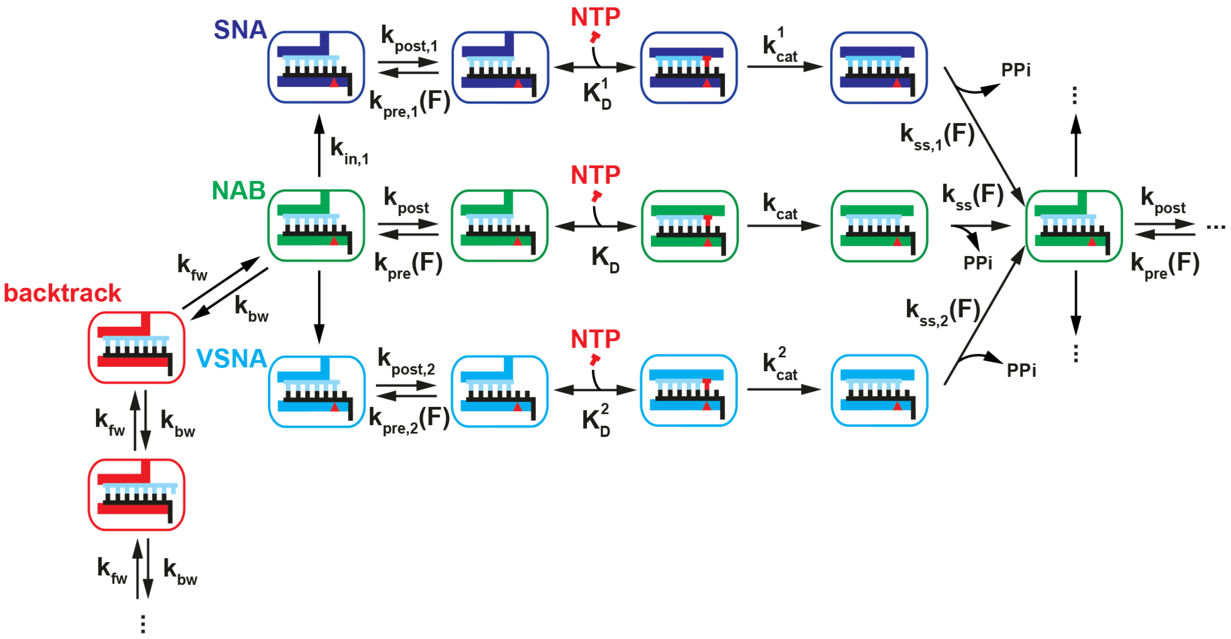

**Figure S9: A detailed mechanochemical model of the coronavirus nucleotide addition cycle**. The model presented here is similar to the one presented in **Fig. 6A**, including the different rates used in our analysis, and their respective dependence on either force or NTP concentration. We also provided a schematic representation of the polymerase active site, which is either open (until nucleotide binding and after pyrophosphate (PPi) release and "reset" of the polymerase active site) or closed. The polymerase elongates a product strand (light blue) by reading the product strand. The red triangle indicates the register of the template strand (black) with the active site. The upstream part of the dsRNA product is not represented. The backtrack kinetics are also described with forward and backward rates, $k_{fw}$ and $k_{bw}$, for the polymerase to jump either one base forward or one base backward, respectively.

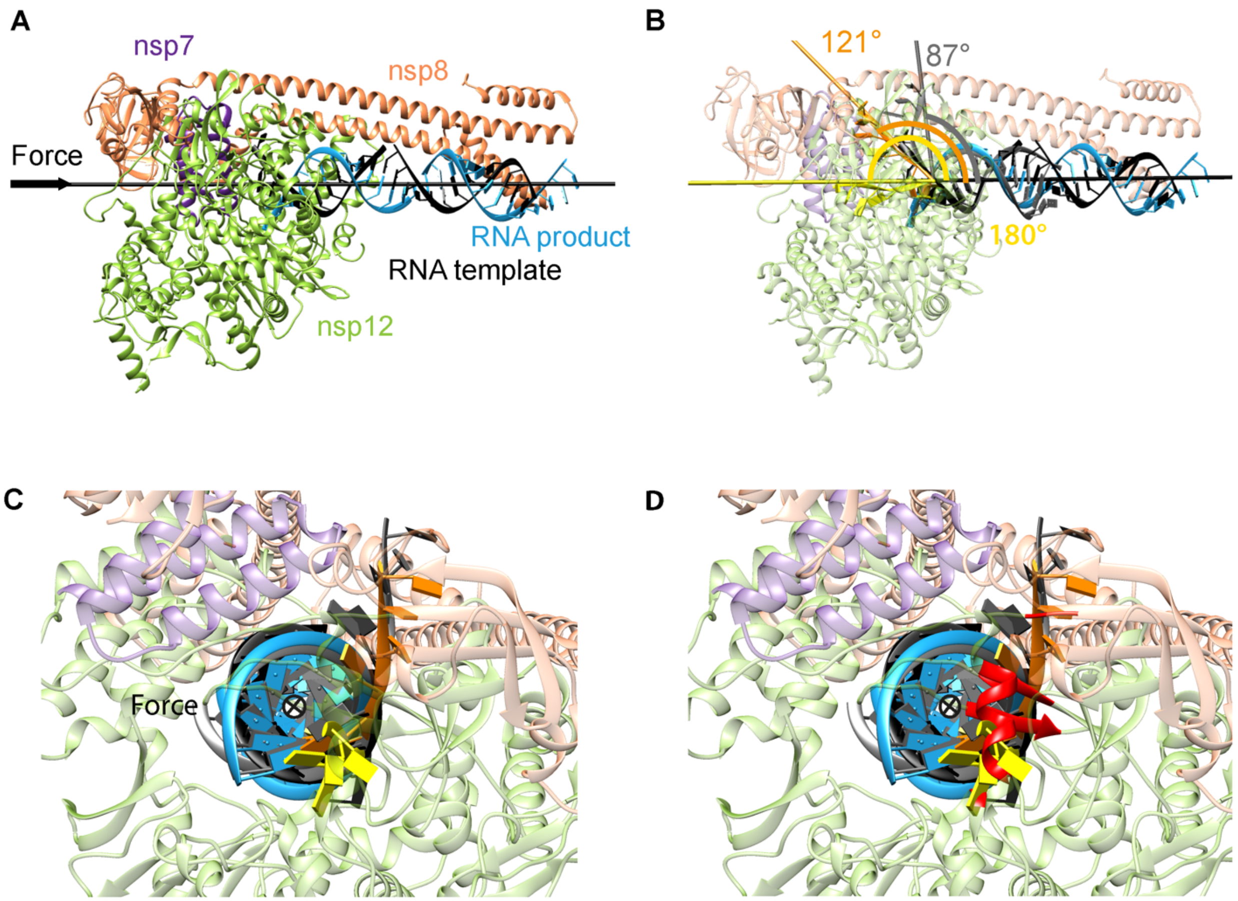

**Figure S10: Structural model SARS-CoV-2 polymerase downstream template strand position (A)** Cryo-EM reconstruction of SARS-CoV-2 polymerase-RNA complex (pdb: 6yyt) as published in Ref. (*14*). The described color code is used throughout. The black axes represent the axe along which the force is applied on the RNA using the magnetic tweezers. **(B)** The (grey) ssRNA is the template RNA entering the RdRp poliovirus as resolved in Ref. (*15*) (pdb: 3ol6, polymerase not shown) was overlapped with the SARS-CoV-2 RNA product using UCSF Chimera 1.15. The angle between the force axe (black) and an axe through the backbone (grey) of the ssRNA template is 87°. Bending of the ssRNA template in direction of the acting force increases this angle up to 121º (orange) until the template RNA clashes with nsp12 residues, which may be skirted to align with the force axis (yellow). **(C)** 90°C rotation of (B), **(D)** with the above mentioned nsp12 residues clashing with the template RNA in red (amino acids M542-L544).

#### Table S1

| nsp12 conc. (μM) | nsp7 conc. (μM) | nsp8 conc. (μM) | NTP conc. (μM) | Force (pN) | Product length | | Total replication time | Dwell time distribution | | | | | | | | Figures |
|---|---|---|---|---|---|---|---|---|---|---|---|---|---|---|---|---|
| | | | | | # traces | (mean ± std) nt | (median ± std) s | # traces | # dwell times | (nucleotide addition rate ± std) 1/s | (pause 1 exit rate ± std) 1/s | (pause 2 exit rate ± std) 1/s | pause 1 probability ± std | pause 2 probability ± std | long-lived (backtrack) pause probability ± std | |
| 0,1 | 0,9 | 0,9 | 500 A/C/G/ UTP | 35 | 58 | 874 ± 36 | 28.0 ± 1.0 | 79 | 6264 | 67.0 ± 1.2 | 4.66 ± 0.20 | 1.65 ± 0.22 | 0.085 ± 0.005 | 0.0052 ± 0.0017 | 0.0004 ± 0.0001 | 2B-E, S2A |
| 0,2 | 1,8 | 1,8 | | | 102 | 867 ± 29 | 26.9 ± 0.8 | 120 | 9622 | 70.7 ± 0.6 | 4.41 ± 0.11 | 0.97 ± 0.18 | 0.078 ± 0.002 | 0.0024 ± 0.0006 | 0.0006 ± 0.0001 | 2B-E, S2A |
| 0,4 | 3,6 | 3,6 | | | 136 | 929 ± 17 | 33.7 ± 1.2 | 165 | 14125 | 66.9 ± 0.6 | 3.98 ± 0.09 | 1.57 ± 0.13 | 0.097 ± 0.003 | 0.0052 ± 0.0011 | 0.0006 ± 0.0001 | 2A-E, S2AB |
| 0,6 | 1,8 | 1,8 | 500 A/C/G/ UTP | 35 | 153 | 924 ± 15 | _ | 101 | 8176 | 66.8 ± 0.7 | 4.2 ± 0.1 | 1.25 ± 0.22 | 0.088 ± 0.003 | 0.0021 ± 0.0009 | 0.0007 ± 0.0002 | 2ABDE, S2B |
| 0,6 | 1,8 | 1,8 | 500 A/C/G/ UTP | 20 | 93 | 880 ± 18 | 18.9 ± 0.5 | 178 | 12152 | 85.2 ± 0.6 | 4.11 ± 0.08 | 1.33 ± 0.28 | 0.060 ± 0.001 | 0.0009 ± 0.0004 | 0.0005 ± 0.0001 | 3A-F, S3A-K |
| | | | | 23 | 116 | 863 ± 17 | 21.2 ± 0.5 | 172 | 13429 | 78.7 ± 0.4 | 4.51 ± 0.10 | 1.48 ± 0.37 | 0.060 ± 0.002 | 0.0009 ± 0.0004 | 0.0011 ± 0.0001 | 3B-F, S3AK |
| | | | | 25 | 93 | 924 ± 12 | 21.1 ± 0.5 | 191 | 16262 | 77.7 ± 0.5 | 5.47 ± 0.13 | 1.74 ± 0.32 | 0.062 ± 0.002 | 0.0018 ± 0.0007 | 0.0008 ± 0.0001 | 1B-D, 3B-F, 4BDE, S3BK, S4A-C |
| | | | | 27 | 92 | 879 ± 20 | 23.8 ± 0.8 | 101 | 8203 | 72.6 ± 0.6 | 4.30 ± 0.09 | 1.55 ± 0.25 | 0.064 ± 0.002 | 0.0013 ± 0.0006 | 0.0009 ± 0.0001 | 3B-F, S3CK |
| | | | | 30 | 168 | 893 ± 18 | 24.6 ± 0.6 | 152 | 12686 | 67.6 ± 0.4 | 4.30 ± 0.09 | 1.80 ± 0.28 | 0.067 ± 0.002 | 0.0013 ± 0.0006 | 0.0006 ± 0.0001 | 3B-F, S3DK |
| | | | | 35 | 269 | 940 ± 13 | 24.7 ± 0.5 | 343 | 29458 | 68.5 ± 0.5 | 4.51 ± 0.11 | 1.63 ± 0.22 | 0.077 ± 0.002 | 0.0034 ± 0.0011 | 0.0008 ± 0.0001 | 3B-F, S3EK, S5B-F |
| | | | | 40 | 117 | 966 ± 18 | 28.5 ± 0.5 | 158 | 14024 | 69.8 ± 0.5 | 4.15 ± 0.09 | 1.16 ± 0.20 | 0.075 ± 0.002 | 0.0023 ± 0.0007 | 0.0004 ± 0.0001 | 3B-F, S3FK |
| | | | | 45 | 138 | 894 ± 24 | 37.7 ± 1.4 | 184 | 15907 | 67.3 ± 0.6 | 3.46 ± 0.06 | 1.02 ± 0.09 | 0.099 ± 0.002 | 0.0041 ± 0.0007 | 0.0007 ± 0.0001 | 3B-F, S3GK |
| | | | | 50 | 86 | 844 ± 25 | 61.9 ± 3.2 | 102 | 7515 | 70.3 ± 1.4 | 2.80 ± 0.09 * | 0.65 ± 0.12 | 0.382 ± 0.054 | 0.0117 ± 0.0034 | 0.0017 ± 0.0005 | 3B-F, S3HK |
| | | | | 55 | 86 | 826 ± 29 | 71.0 ± 4.0 | 97 | 7515 | 67.3 ± 1.8 | 2.38 ± 0.04 * | 0.53 ± 0.05 | 0.356 ± 0.040 | 0.0124 ± 0.0019 | 0.0018 ± 0.0003 | 3B-F, S3IK |
| | | | | 60 | 71 | 806 ± 32 | 82.6 ± 8.2 | 92 | 6822 | 62.8 ± 1.4 | 2.23 ± 0.05 * | 0.54 ± 0.04 | 0.332 ± 0.031 | 0.0140 ± 0.0022 | 0.0023 ± 0.0004 | 3A-F, S3JK |
| 0,6 | 1,8 | 1,8 | 1000 | 35 | 132 | 933 ± 19 | 25.2 ± 0.8 | 200 | 17537 | 68.0 ± 0.5 | 4.97 ± 0.11 | 1.45 ± 0.19 | 0.067 ± 0.002 | 0.0015 ± 0.0005 | 0.0005 ± 0.0001 | S5A-F |
| | | | 500 | | 269 | 940 ± 13 | 24.7 ± 0.5 | 343 | 29458 | 68.5 ± 0.5 | 4.51 ± 0.11 | 1.63 ± 0.22 | 0.077 ± 0.002 | 0.0034 ± 0.0011 | 0.0008 ± 0.0001 | 3B-F, S3EK, S5B-F |
| | | | 300 | | 143 | 957 ± 16 | 33.2 ± 1.4 | 178 | 15088 | 63.2 ± 0.6 | 4.17 ± 0.15 | 1.42 ± 0.16 | 0.088 ± 0.003 | 0.0082 ± 0.0020 | 0.0006 ± 0.0001 | S5B-F |
| | | | 100 | | 172 | 911 ± 18 | 53.0 ± 1.2 | 183 | 15191 | 65.6 ± 0.8 | 3.01 ± 0.04 * | 0.78 ± 0.05 | 0.189 ± 0.006 | 0.0061 ± 0.0008 | 0.0007 ± 0.0001 | S5B-F |
| | | | 50 | | 133 | 985 ± 12 | 98.0 ± 2.5 | 155 | 13770 | 66.1 ± 1.3 | 1.57 ± 0.02 * | 0.27 ± 0.02 | 0.315 ± 0.015 | 0.0038 ± 0.0005 | 0.0008 ± 0.0002 | S5B-F |
| | | | 20 | | 135 | 922 ± 18 | 262.8 ± 9.5 | 155 | 13452 | 44.7 ± 5.8 | 0.59 ± 0.01 * | 0.01 ± 0.11 | 0.343 ± 0.020 | 0.0050 ± 0.0007 | 0.0020 ± 0.0003 | S5B-F |
| | | | 10 | | 104 | 909 ± 23 | 367.1 ± 14.0 | 123 | 10473 | 25.8 ± 5.7 | 0.39 ± 0.01 * | 0.08 ± 0.01 | 0.304 ± 0.023 | 0.0058 ± 0.0010 | 0.0014 ± 0.0003 | S5A-F |
| 0,6 | 1,8 | 1,8 | 1000 | 25 | 82 | 916 ± 16 | 21.4 ± 0.5 | 124 | 10442 | 78.1 ± 0.5 | 4.07 ± 0.10 | 1.51 ± 0.33 | 0.055 ± 0.002 | 0.0008 ± 0.0005 | 0.0008 ± 0.0001 | 4A-E, S4AC |
| | | | 500 | | 93 | 924 ± 12 | 21.1 ± 0.5 | 191 | 16262 | 77.7 ± 0.5 | 5.47 ± 0.13 | 1.74 ± 0.32 | 0.062 ± 0.002 | 0.0018 ± 0.0007 | 0.0008 ± 0.0001 | 1B-D, 3B-F, 4B-E, S3BK, S4A-C |
| | | | 300 | | 133 | 889 ± 17 | 25.3 ± 0.6 | 194 | 15612 | 74.6 ± 0.6 | 4.55 ± 0.08 | 1.65 ± 0.23 | 0.080 ± 0.002 | 0.0010 ± 0.0003 | 0.0006 ± 0.0001 | 4B-E, S4A-C |
| | | | 100 | | 305 | 873 ± 12 | 40.9 ± 0.7 | 348 | 28172 | 73.9 ± 0.8 | 2.83 ± 0.03 * | 0.45 ± 0.09 | 0.144 ± 0.003 | 0.0013 ± 0.0003 | 0.0006 ± 0.0001 | 4B-E, S4A-C |
| | | | 50 | | 174 | 875 ± 14 | 69.9 ± 2.0 | 187 | 15148 | 75.7 ± 1.3 | 1.82 ± 0.02 * | 0.35 ± 0.07 | 0.229 ± 0.007 | 0.0014 ± 0.0003 | 0.0018 ± 0.0003 | 4B-E, S4A-C |
| | | | 20 | | 113 | 894 ± 18 | 164.8 ± 6.6 | 123 | 10449 | 76.6 ± 2.0 | 0.82 ± 0.01 * | 0.10 ± 0.02 | 0.344 ± 0.010 | 0.0018 ± 0.0003 | 0.0014 ± 0.0003 | 4B-E, S4A-C |
| | | | 10 | | 137 | 859 ± 20 | 282.7 ± 8.6 | 146 | 11820 | 76.0 ± 1.4 | 0.44 ± 0.01 * | 0.06 ± 0.01 | 0.355 ± 0.008 | 0.0016 ± 0.0004 | 0.0017 ± 0.0004 | 4A-E, S4AC |
| 0,2 | 1,8 | 1,8 | 500 A/C/G/ UTP | 9 | 14 | 419 ± 29 | 123.5 ± 26.8 | 45 | 1586 | 41.9 ± 1.5 | 0.80 ± 0.08 | 0.18 ± 0.03 | 0.098 ± 0.007 | 0.0190 ± 0.0049 | 0.0107 ± 0.0025 | 5B, 5E-H, S6C |
| | | | | 12 | 63 | 409 ± 11 | 51.7 ± 3.0 | 83 | 2953 | 44.8 ± 0.9 | 1.02 ± 0.05 | 0.18 ± 0.02 | 0.091 ± 0.004 | 0.0102 ± 0.0015 | 0.0029 ± 0.0008 | 5E-H, S6C |
| | | | | 15 | 45 | 412 ± 15 | 38.6 ± 3.7 | 98 | 3419 | 39.5 ± 0.7 | 2.18 ± 0.41 | 0.49 ± 0.16 | 0.049 ± 0.005 | 0.0135 ± 0.0050 | 0.0063 ± 0.0014 | 5E-H, S6C |
| | | | | 18 | 65 | 426 ± 11 | 30.4 ± 2.8 | 97 | 3886 | 45.3 ± 1.2 | 3.01 ± 1.11 | 0.73 ± 0.27 | 0.053 ± 0.009 | 0.0202 ± 0.0109 | 0.0036 ± 0.0012 | 5E-H, S6C |
| | | | | 20 | 74 | 425 ± 8 | 13.8 ± 0.7 | 114 | 4295 | 52.4 ± 1.1 | 5.01 ± 0.44 | 1.17 ± 0.14 | 0.034 ± 0.005 | 0.0138 ± 0.0020 | 0.0014 ± 0.0004 | 5B, 5E-H, S6C |

* rates corrected
